# Aberrant RNA methylation triggers recruitment of an alkylation repair complex

**DOI:** 10.1101/2020.08.28.271874

**Authors:** Joshua R. Brickner, Ning Tsao, Rebecca Rodell, Clement Oyeniran, Valentina Lukinović, Albino Bacolla, Lisheng Zhang, Jennifer M. Soll, Alexandre G. Casanova, Adit Ganguly, Chuan He, John A. Tainer, Nicolas Reynoird, Nima Mosammaparast

**Author notes:** These authors contributed equally to this work (Joshua R. Brickner, Ning Tsao). Correspondence to: Nima Mosammaparast Department of Pathology & Immunology Washington University School of Medicine 4940 Parkview Place, CSRB Room 7750 St. Louis MO, 63110.

## Abstract

A critical question in genome stability is the nature of the chemical damage responsible for repair activation. We previously reported a novel pathway specifically activated during alkylation damage in human cells, where the E3 ubiquitin ligase RNF113A mediates the recruitment of the ASCC repair complex. Yet the mechanistic basis for the alkylation damage selectivity of this pathway remains unclear. Here, we demonstrate that RNA but not DNA alkylation is the initiating signal for this process. Aberrantly methylated RNA is sufficient to recruit ASCC, while an RNA dealkylase suppresses ASCC recruitment during chemical alkylation. This aberrant RNA methylation causes transcriptional repression in a manner dependent on the ASCC complex. We show that an alkylated pre-mRNA, or an RNA containing a single damaged base, is sufficient to activate RNF113A E3 activity in a phosphorylation-dependent manner. Together, our work identifies an unexpected role for RNA damage in eliciting a DNA repair response, and suggests that RNA may serve as the “canary in the coal mine” for sensing alkylation damage.

## Introduction

The vital importance of genome maintenance is underscored by the evolution of multiple mechanisms to repair specific types of damaged DNA. Repair of alkylated DNA is particularly critical in the context of cancer, since alkylation chemotherapy is used to treat many types of tumors (Fu et al., 2012; Soll et al., 2017). In addition, alkylation can be caused by endogenous sources, such as the universal methyl donor S-adenosyl-methionine, as well as nitrosamines or bile acids (Garcia-Santos Mdel et al., 2001; Rydberg and Lindahl, 1982). These agents react with DNA bases on either nitrogen or oxygen atoms via an S_N_1 or S_N_2 nucleophilic substitution reaction. Depending on the type of modification, alkylated bases can disrupt Watson-Crick base pairing, inhibiting DNA polymerases or inducing misincorporation during replication (Fu et al., 2012; Soll et al., 2017). Three major pathways are responsible for the repair of alkylated DNA, including base excision repair (BER) as well as the AlkB family of oxidative demethylases, both of which repair distinct *N*-linked lesions, and methylguanine methyltransferase (MGMT), which reverts *O*-linked lesions (Brickner et al., 2019; Drablos et al., 2004; Fu et al., 2012). Additionally, nucleotide excision repair (NER) and the Fanconi anemia (FA) pathway can remove bulky alkylated adducts or interstrand crosslinks induced by bifunctional alkylators, respectively (Kim and D’Andrea, 2012; Spivak, 2015). Clear evidence exists linking each of these pathways to hypersensitivity or resistance to alkylation damage responses in tumors (Butler et al., 2020; Fu et al., 2012; Soll et al., 2017). Yet despite their vast biochemical and functional characterization, how any specific alkylation repair pathway is selectively activated and deployed in human cells, or how they are coordinated with DNA-templated processes such as transcription, remains largely elusive.

Alkylating agents cause lesions not only in DNA, but also in RNA (Fedeles et al., 2015; Vagbo et al., 2013). In fact, treatment of *E. coli* with the alkylating agent methyl methyanesulphonate (MMS) resulted in 10-fold more abundant alkylated RNA than alkylated DNA (Vagbo et al., 2013). Methylated bases induced by alkylation, such as *N*1-methyladenine (m1A), *N*3-methylcytosine (m3C) and *N*7-methylguanine (m7G) are also common RNA modifications in the cell (Shi et al., 2019; Thapar et al., 2019). In the cytoplasm, m1A and m7G have been shown to regulate translation efficiency as well as maintaining RNA stability (Li et al., 2017; Lin et al., 2018; Safra et al., 2017; Zhang et al., 2019). The overlap between these marks and damage lesions implies that the cellular effects of alkylation may also extend to alter the physiological functions regulated by these RNA methylation marks. Alkylated mRNAs likely induce various RNA degradation pathways, particularly the ribosome-quality-control pathway (RQC) (Yan et al., 2019). Intriguingly, RNA dealkylation mechanisms exist, yet compared to the well-studied repair pathways for DNA, the repair systems responsible for RNA repair are poorly understood. Some viral AlkB proteins and *E. coli* AlkB are capable of repairing m1A and m3C in both single-stranded DNA and RNA (Aas et al., 2003; van den Born et al., 2008). Clear evidence for RNA dealkylation activity has also been demonstrated for a number of the human AlkB proteins, including ALKBH1, ALKBH3, and ALKBH8 (Aas et al., 2003; Li et al., 2017; Liu et al., 2016; Ueda et al., 2017; van den Born et al., 2011). Emerging from these studies is the notion that alkylated RNA is likely to affect cellular physiology and organisms appear to have evolved pathways to degrade or repair damaged RNA.

We previously showed that ALKBH3 is coupled to the ASCC helicase complex, which in turn is specifically recruited during alkylation damage to nuclear speckle bodies (Brickner et al., 2017). These regions are euchromatic centers associated with RNA polymerase II transcript processing. The recruitment of this repair complex requires K63-linked ubiquitination mediated by the RNF113A E3 ubiquitin ligase and recognition of these ubiquitin chain by the ASCC2 accessory subunit. In a similar ubiquitin-dependent manner, the ASCC complex is also recruited to mediate RQC in the cytosol (Juszkiewicz et al., 2020; Matsuo et al., 2020). This ubiquitin-dependent mechanism is reminiscent of other more established damage-induced signaling pathways, such as DNA double-stranded break repair, which activates non-canonical ubiquitination to recruit damage effector complexes (Schwertman et al., 2016). Yet the molecular basis for the alkylation damage specificity in the RNF113A-ASCC-ALKBH3 pathway remains unclear.

Here, we demonstrate that aberrant nuclear RNA methylation is both necessary and sufficient to activate this pathway normally induced upon chemical alkylation. The recruitment of ASCC mediates transcriptional repression, both globally and locally, suggesting a role in ridding cells of alkylated nascent transcripts. We find that RNF113A E3 ligase activity is induced specifically during methylation damage, but not with other types of genotoxic agents. Mechanistically, we reveal that the RNF113A E3 ligase complex mediates recognition of alkylated RNAs and that its E3 activity is regulated through a phosphorylation-dependent mechanism. Our findings uncover a novel role for aberrantly methylated RNA in initiating the cellular response to alkylation via the activation of the RNF113A-ASCC pathway.

## Results

### RNA alkylation is necessary and sufficient to mediate ASCC recruitment

We previously reported the discovery of a nuclear ubiquitin-dependent signaling pathway that is specifically required for recruiting the ASCC complex to mediate alkylation damage response (Brickner et al., 2017). Although the mechanistic basis for this alkylation damage selectivity has remained unclear, we reasoned that the specificity of this pathway towards alkylation may be due to modification of RNA, consistent with the fact that RNA is more easily modified by alkylating agents relative to DNA and as such may serve as early sensor for alkylation stress (Drablos et al., 2004; Vagbo et al., 2013). To test whether RNA methylation was necessary for ASCC recruitment, we wished to use an RNA-specific repair enzyme to counter the alkylation-mediated modification of RNA without affecting DNA alkylation. Therefore, we cloned and characterized an AlkB demethylase from blueberry scorch virus (BsV), an RNA virus (van den Born et al., 2008). Consistent with previous data, the BsV-AlkB dealkylase was found to be active on m1A-modified RNA olignonucleotide but not on an identical DNA substrate *in vitro*, as assessed by quantitative LC-MS/MS (Figure 1A; Supplemental Figure S1A-S1C). Having established its selectivity for RNA, we next expressed BsV-AlkB as an NLS-fusion in U2OS cells, which targeted it to the nucleus (Supplemental Figure S1D). Strikingly, BsV-AlkB-NLS expression significantly reduced HA-ASCC2 foci formation to background levels following MMS-induced alkylation damage (Figure 1B-C). The enzymatic activity of BsV-AlkB was important for foci suppression, as the H156A catalytic mutation was less capable of countering ASCC2 foci formation, suggesting that RNA alkylation is important for MMS-induced ASCC2 foci formation.

**Figure 1.**
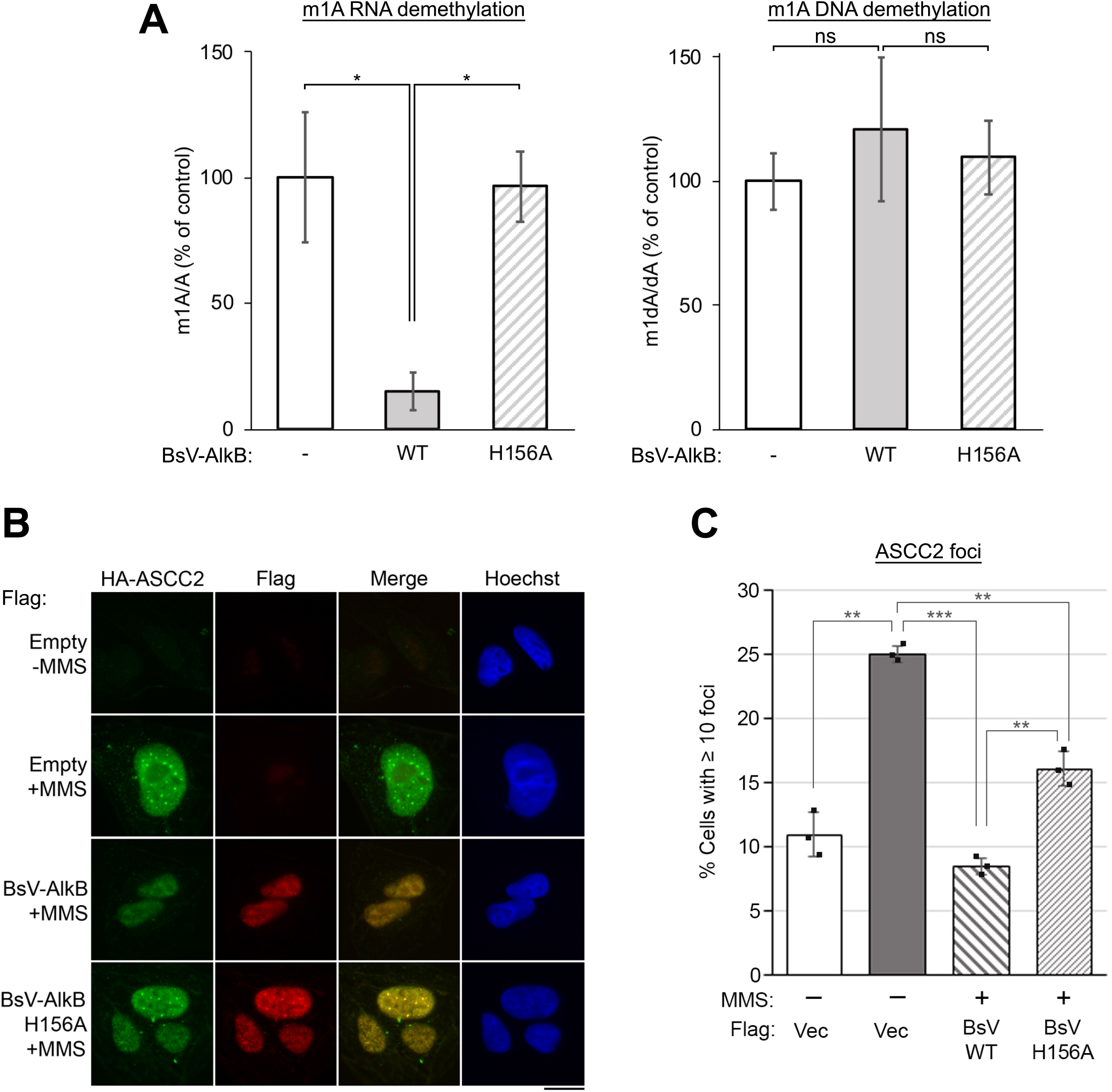
RNA alkylation is necessary for ASCC recruitment to nuclear foci during alkylation damage. **(A)** *In vitro* demethylation assays on identical sequences of an m1A-containing RNA oligonucleotide (left) or DNA oligonucleotide (right) with wildtype or catalytically inactive (H156A) recombinant BsV-AlkB protein. Reactions were quantified by LC-MS/MS and shown as a percent of no protein control (n=4 to 5 independent replicates for each reaction; error bars indicate ± S.D. of the mean; * = *p* < 0.05).**(B)** U2OS cells expressing HA-ASCC2, as well as Flag empty vector, or the indicated versions of Flag BsV-AlkB-NLS were treated with MMS (500 μM) for six hours, then processed for immunofluorescence after Triton X-100 extraction using the indicated antibodies. Scale bar, 10 μm. **(C)** Quantification of **(B)**. (n=3 replicates; error bars indicate ± S.D. of the mean; two-tailed *t*-test; ** = *p* < 0.001, *** = *p* <0.0001).

To test if RNA methylation was sufficient to recruit the ASCC complex, we turned to RNA methyltransferases that produce so-called “intentional” methylated bases on rRNA and tRNA, which are identical to damage marks formed by reactions of RNA with chemical agents. We cloned the METTL8 methyltransferase, which has been shown to produce 3-methylcytosine (m3C) on various RNAs, including mRNA (Xu et al., 2017). While the function of this modification on mRNA is unclear, it is one of the major modifications produced on RNA by MMS and other methylating agents and is also a substrate for the AlkB family of repair enzymes (Aas et al., 2003; Drablos et al., 2004; Ougland et al., 2004). We purified METTL8 and various catalytic mutants from 293T cells (Supplemental Figure S2A and S2C). As expected, using LC-MS/MS METTL8 was found to be active only on a ssRNA substrate *in vitro*; no methyltransferase activity on the same ssDNA substrate was detected (Figure 2A-2B). We next cloned and expressed a human METTL8-NLS fusion, which was targeted to the nucleus, particularly to the nucleolus (Supplemental Figure S2A-S2B). More importantly and consistent with our model that RNA damage plays an important role in ASCC recruitment, ectopic expression of wildtype METTL8-NLS induced recruitment of the ASCC3 helicase to nucleolar regions, which coincided with localization of the active methyltransferase (Figure 2C-2D). Conversely, two catalytic mutations targeting the putative SAM-binding domain of METTL8 (D’Silva et al., 2011; Noma et al., 2011) (G204A/G206A and D230A; Supplemental Figure S2A-S2C) failed to recruit ASCC3 to nucleoli (Figure 2C-2D and Supplemental Figure S2D). LC-MS/MS confirmed that these two mutants were deficient for RNA methyltransferase activity (Figure 2A-2B). We noted that the mutant METTL8-NLS proteins did not localize to nucleoli as robustly as their wildtype counterpart (Supplemental Figure S2B), which could alternatively explain the impairment of ASCC3 nucleolar recruitment. To clarify this question, we utilized an integrated single locus system where we could target this methyltransferase to an MS2 gene reporter (Janicki et al., 2004; Shanbhag et al., 2010). In this system, fusing a degron-tagged METTL8-NLS to mCherry-LacI allowed for targeted recruitment of this RNA methyltransferase to the locus upon induction with the Shield1 ligand (Figure 2E). Although the WT fusion protein still partially localized to nucleoli, we found that METTL8 similarly induced ASCC3 recruitment to this locus, which in contrast, was markedly attenuated with the METTL8 G204A/G206A catalytic mutant (Figure 2F-G). Taken together, our results support the notion that aberrant RNA methylation is both necessary and sufficient to recruit the ASCC complex.

**Figure 2.**
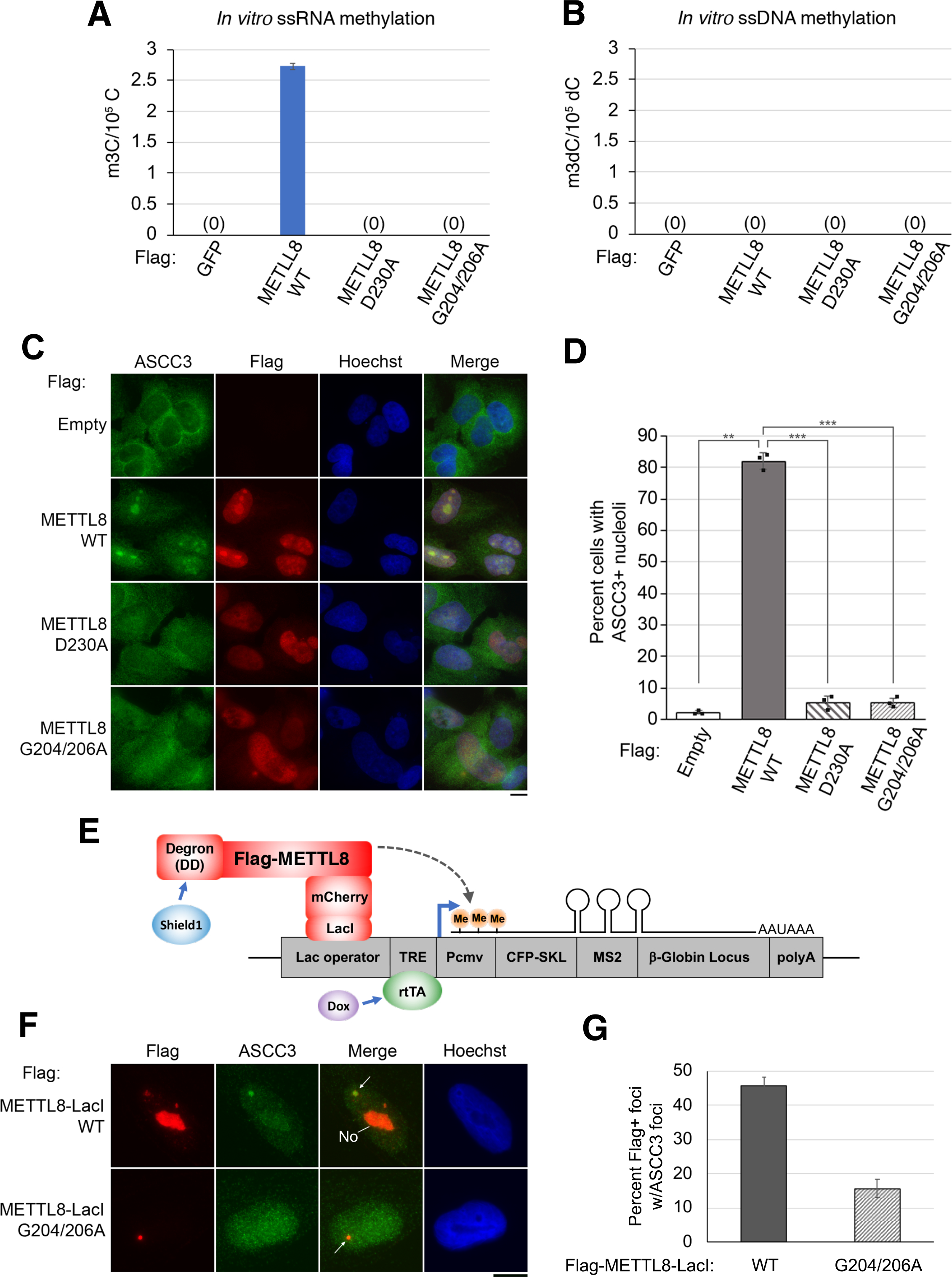
RNA alkylation is sufficient to mediate ASCC recruitment. **(A)** and **(B)** *In vitro* methyltransferase assays were performed using the four indicated Flag-tagged proteins with an RNA **(A)** or DNA **(B)** oligonucleotide substrate. Products were digested to nucleosides and quantified using LC-MS/MS. N = 2 independent replicates and error bars indicate ±S.D. of the mean. **(C)** U2OS cells expressing the empty Flag vector, or the indicated Flag-METTL8-NLS vectors, were processed for immunofluorescence using the indicated antibodies. **(D)** Quantification of **(C)** (n=3 replicates; error bars indicate ± S.D. of the mean; ** = *p* < 0.001, *** = *p* < 0.0001). **(E)** Schematic of locus reporter system to target RNA methylation using the DD-Flag-METTL8-mCherry-LacI construct. **(F)** METTL8-catalyzed ASCC3 recruitment to the targeted locus visualized by immunofluorescence as shown. ‘No’ indicates nucleolus, arrow indicates reporter locus. **(G)** Quantification of **(F)**. n=2 replicates; error bars indicate ± S.D. of the mean.

### ASCC-mediated transcriptional repression in response to aberrant RNA methylation

Previous work using various genotoxins has revealed that DNA damage generally elicits a repressive transcriptional response, either locally or globally (Purman et al., 2019; Williamson et al., 2017). We reasoned that alkylation damage may also result in transcriptional repression, potentially due to damage to RNA. To this end, RNA-Seq analysis revealed that while global gene repression is not observed in response to MMS, a greater number of genes were downregulated than upregulated (Figure 3A and Supplemental Figure S3A-S3B). Of note, RNA-binding factors and the unfolded protein response were enriched in the upregulated gene set, suggesting these to be a component of the cellular response to alkylation (Supplemental Figure S3A-S3B). We found that many of the genes repressed upon alkylation are at least partially derepressed in ASCC3 KO cells, implicating a role for ASCC3 in the suppression of these genes during such damage (Figure 3B-3C). ASCC3 did not appear to have a global role in transcriptional repression without MMS treatment (Supplemental Figure S3C). Since ASCC2 is a stoichiometric partner of ASCC3 and mediates ubiquitin-dependent targeting of ASCC3 (Brickner et al., 2017), we also analyzed the transcriptional response in ASCC2 deficient cells. Again, we found that loss of ASCC2 results in partial derepression of these same genes (Figure 3C and Supplemental Figure S3D-S3E). Taken together, these data support the view that the ASCC complex is responsible for transcriptional repression of certain genes during alkylation.

**Figure 3.**
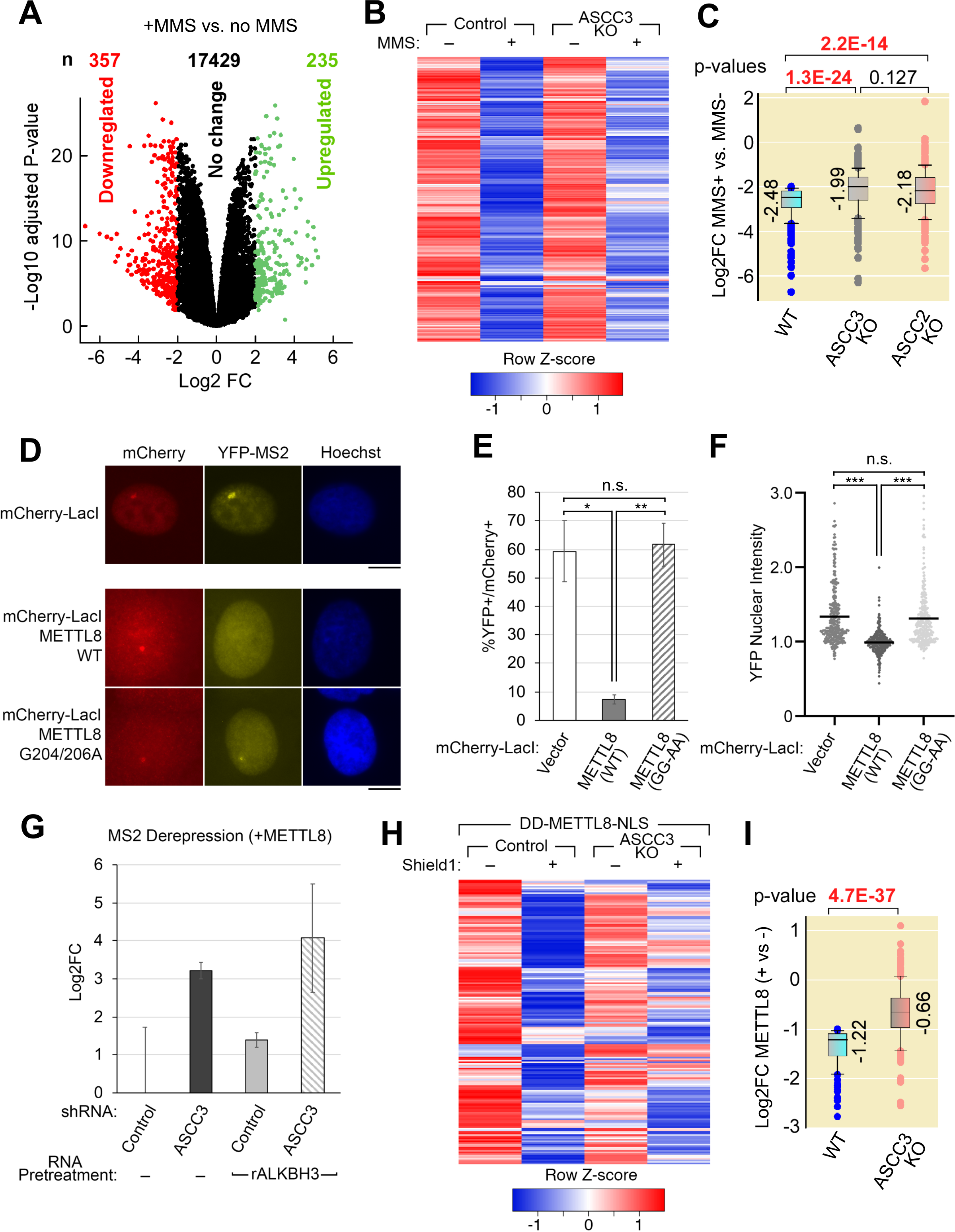
ASCC mediates transcriptional repression in response to aberrant RNA methylation. **(A)** RNA-Seq transcriptome analysis of WT U2OS cells with and without MMS treatment (n=3 biological replicates per condition). Genes upregulated or downregulated more than ±2 log_2_ fold change (FC) upon MMS treatment are highlighted in green and red, respectively. **(B)** Heatmap comparing MMS-downregulated genes in WT and ASCC3 KO cells (n=3 biological replicates per condition). **(C)** Box plot of downregulated genes upon MMS treatment in WT cells in comparison to ASCC3 and ASCC2 KO cells. **(D)** METTL8-mediated gene repression as visualized by the MS2-YFP reporter system. The mCherry-LacI vector alone is shown separately due to a different exposure time used solely for the mCherry-LacI control protein. **(E)** Quantification of MS2-YFP-positive/mCherry-positive foci from **(D)** (n=3 independent replicates; error bars indicate ± S.D. of the mean; * *p* < 0.05, ** p < 0.01, n.s. not significant). **(F)** Quantification of nuclear foci intensities from **(D)** (n=3 independent replicates; error bars indicated ± S.D. of the mean; *** *p* < 0.001, n.s. not significant). **(G)** MS2 derepression as determined by quantitative RT-PCR from control versus ASCC3 knockdown cells, using β-actin as the housekeeping control. Where indicated, the RNA was treated with purified ALKBH3 prior qRT-PCR (n=3 biological replicates; error bars indicated ± S.D. of the mean). **(H)** Heatmap comparing DD-METTL8-NLS-mediated downregulated genes in WT and ASCC3 KO cells (n=3 biological replicates per condition). **(I)** Box plot of downregulated genes upon DD-METTL8-NLS induction with Shield1 ligand in WT cells in comparison to ASCC3 KO cells.

To test whether the transcriptional response during alkylation could be due to aberrant RNA methylation of nascent transcripts, we used the reporter system from Figure 2 to assess whether METTL8 recruitment would result in repressed transcription of the MS2 reporter. Indeed, targeting METTL8 to the single locus reporter significantly repressed transcription (Figure 3D-3F and Supplemental Figure S4A). This repression of nascent transcription depended upon METTL8 methyltransferase activity (Figure 3D-3F). We verified that deposition of m3C on the nascent transcript did not inhibit its quantitation by qRT-PCR, since our results were not significantly altered after demethylation of the purified RNA with recombinant ALKBH3, which preferentially removes m3C from RNA(Aas et al., 2003) (Supplemental Figure S4B). To test whether loss of ASCC3 would alter the expression of nascent transcription from this single locus reporter, we performed qRT-PCR upon shRNA-mediated knockdown of ASCC3 (Figure 3G and Supplemental Figure S4C). As with the transcriptional response to MMS, loss of ASCC3 significantly derepressed this target locus. To determine whether this represented a general phenomenon in response to global RNA methylation within the nucleus, we expressed METTL8-NLS in WT and ASCC3 KO cells and analyzed transcription by RNA-Seq (Figure 2H and Supplemental Figure S4D). While the overall transcriptional changes were relatively moderate in response to METTL8-NLS expression in comparison to MMS treatment (Supplemental Figure S4D), we again found that loss of ASCC3 caused derepression of genes that were downregulated by nuclear METTL8 (Figure 3H-3I). Altogether, our data strongly suggests that the ASCC complex is recruited to sites harboring aberrant RNA methylation to downregulate their expression.

### Selective activation of RNF113A E3 ligase upon alkylation damage

We wished to determine the molecular basis of the RNA alkylation-mediated recruitment of the ASCC complex. A key factor upstream of the recruitment of this complex is the E3 ubiquitin ligase RNF113A (Brickner et al., 2017), which we reasoned could be selectively activated during alkylation. Many E3 ubiquitin ligases undergo autoubiquitination when activated (Zheng and Shabek, 2017), and indeed, we found that RNF113A is ubiquitinated in cells upon MMS treatment (Figure 4A). Deletion of the RING domain, or a catalytic point mutation (I264A) within this domain (Brickner et al., 2017), significantly reduced RNF113A ubiquitination during MMS treatment, supporting the notion that this represented RNF113A autoubiquitination (Supplemental Figure S5A). Other types of damaging agents, including hydroxyurea, bleomycin, or camptothecin failed to induce RNF113A autoubiquitination (Figure 4A). Using tandem ubiquitin binding element (TUBE) conjugated beads to isolate endogenously ubiquitinated proteins, we found that endogenous RNF113A was also autoubiquitinated in response to MMS but not the other types of damaging agents, despite the fact that DNA damage signaling, as assessed by pH2A.X, was apparent under these conditions (Supplemental Figure S5B-S5C). A significant portion of the ubiquitin linkage associated with RNF113A during MMS treatment appeared to be K63-linked, consistent with previous findings demonstrating that RNF113A functions with the K63-specific E2 conjugating enzyme UBC13 (Brickner et al., 2017) (Supplemental Figure S5D). Equimolar concentrations of the methylating agent methyl iodide (MeI) also induced autoubiqitination of RNF113A but the ethylating agent ethyl methanesulphonate (EMS) was significantly less potent in this assay, suggesting that RNF113A is activated primarily by methylation and not by larger alkyl adducts (Figure 4B). Autoubiquitination of RNF113A was accompanied by increased RNF113A E3 ligase activity, as RNF113A purified from cells treated with MMS was significantly more active in producing ubiquitin chains *in vitro* than RNF113A from untreated cells (Supplemental Figure S5E-F). MMS-induced autoubiquitination was specific to RNF113A, as RNF8 and RNF168, two other E3 ligases involved in DNA damage signaling, were not autoubiquitinated during alkylation (Figure 4C). Consistent with its selective activation during alkylation damage, loss of RNF113A resulted in markedly increased sensitivity to MMS but not camptothecin as determined by a colorimetric survival assay as well as colony formation assay (Figure 4D-4E and Supplemental Figure S5G-I). Altogether, our data suggest that RNF11A is selectively activated under alkylation stress and this activation involves its autoubiquitination.

**Figure 4.**
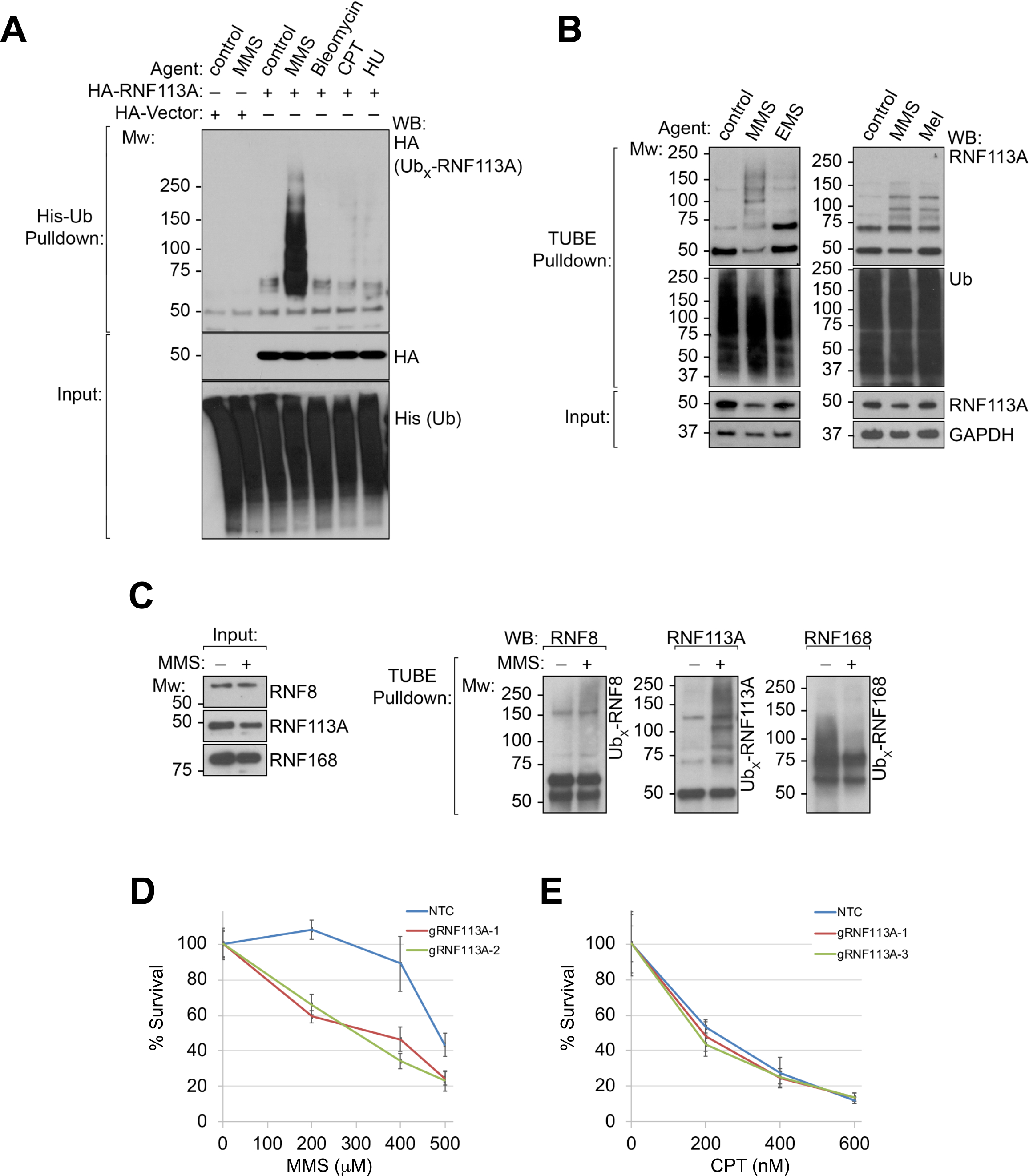
Alkylation damage selectively activates RNF113A E3 ligase activity in cells. **(A)** HeLa cells expressing His-Ub and the indicated HA-vectors were treated with the damaging agents as shown. Whole cell extracts were then used for His-pulldowns using Ni-NTA beads under denaturing conditions. Lysate inputs as well as the pulldown material were analyzed by Western blot using the indicated antibodies (n=3 independent experiments). **(B)** HeLa-S cells were treated with equimolar amounts of MMS versus ethyl methanesulfonate (EMS), or MMS versus methyl iodide (MeI), and cell lysates were used for tandem ubiquitin binding element (TUBE) pulldown using ubiquilin-conjugated beads. Lysate inputs as well as the pulldown material were analyzed by Western blot as shown (n=3 independent experiments). **(C)** TUBE pulldowns were performed as in **(B)** using MMS, then probed for the different E3 ligases by Western blot as shown (right). Input lysates are shown on the left (n=2 independent experiments). **(D)** and **(E)** Control gRNA or RNF113A-specific gRNAs were used for genome editing in HeLa cells. Resistance to MMS **(D)** or camptothecin **(E)** was determined using MTS assay. (n=5 technical replicates for each gRNA/damage condition combination).

### In vitro activation of RNF113A E3 ligase by methylated RNA

The fact that the ASCC complex is recruited by methylation of RNA and that RNF113A is selectively activated by methylating agents in cells strongly suggested that the E3 ligase activity of RNF113A may be activated by methylated RNA. Since we previously found that recombinant RNF113A showed no E3 activity (Brickner et al., 2017), we used RNF113A complex purified from HeLa-S cell nuclear extracts, as above, and tested the ability of methylated RNAs to induce its E3 ligase activity. Due to its association with the spliceosome (Hegele et al., 2012; Shostak et al., 2020), we first used *in vitro* transcribed β-globin pre-mRNA (Movassat et al., 2014) as a potential substrate for inducing RNF113A E3 activity. While the unmodified RNA had a mild stimulatory effect on the E3 activity of RNF113A, we saw a marked increase in the E3 activity when the same RNA was pre-methylated using dimethyl sulphate (DMS; Figure 5A). As expected, this agent induced the common alkylating marks m7G, m1A, and m3C, as determined by LC-MS/MS (Supplemental Figure S6A). The modified RNA alone lacked any apparent E3 activity, suggesting a specific stimulation of RNF113A (Supplemental Figure S6B). We then tested whether synthetic RNA oligonucleotides were capable of increasing the E3 activity of RNF113A. As predicted, a single-stranded RNA oligo containing a single m1A modification, but not its unmodified counterpart, induced the E3 activity of RNF113A (Figure 5B). In contrast, the DNA oligonucleotide version containing m1A in the same position had no apparent effect on inducing increased ubiquitination by the RNF113A complex (Figure 5B).

**Figure 5.**
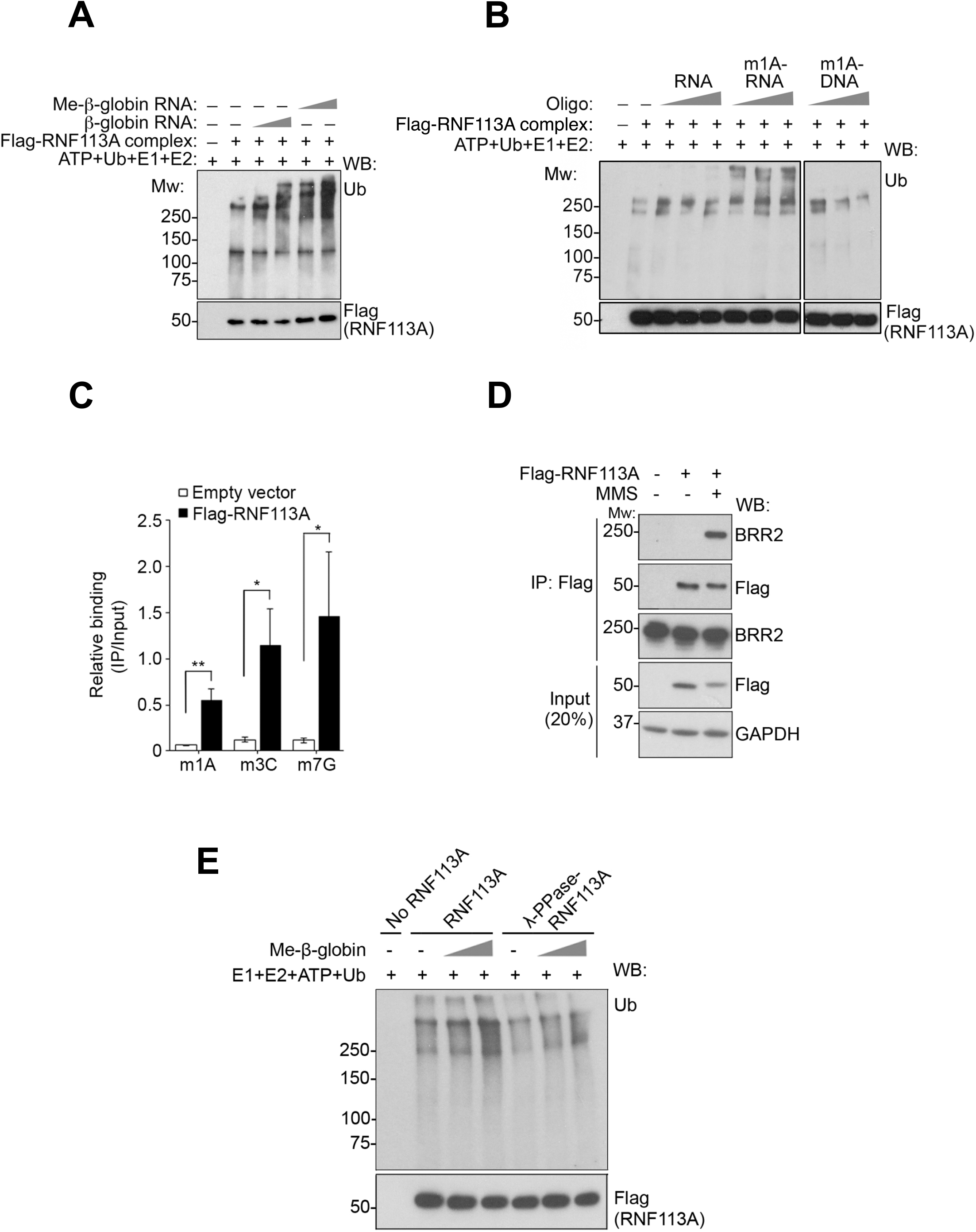
Methylated RNA activates RNF113A *in vitro* in a phosphorylation-dependent manner. **(A)** Flag-RNF113A complex was immunopurified from untreated HeLa-S nuclear lysate, and *in vitro* E3 ligase assays were performed in the presence of the β-globin pre-mRNA (0.1 or 0.3 μM) with or without prior treatment with dimethyl sulfate (DMS) as the methylating agent. The reaction products were analyzed by Western blotting using the antibodies as shown (n=3 independent experiments). **(B)** *In vitro* E3 ligase assays were performed as in **(A)** with the presence of 0.1 μM, 0.2 μM or 0.5 μM single-stranded RNA, m1A-RNA or m1A-DNA oligonucleotides (n=2 independent experiments). Result with the m1A DNA (right) was from the same blot as the m1A RNA. **(C)** HeLa-S cells expressing the empty vector (EV), or Flag-RNF113A were subjected to RNA immunoprecipitation. Purified RNAs were digested and the methylated nucleosides were analyzed by mass spectrometry. The results are shown as the mean ± S.D. (n=3 independent replicates; * *p* < 0.05, ** *p* < 0.01). **(D)** Flag immunoprecipitation was performed from HeLa-S cells expressing empty vector or Flag-RNF113A with or without MMS treatment (5mM for 30 minutes). Bound material, as well as the inputs, were Western blotted with the antibodies as shown (n=3 independent replicates). **(E)** Flag-RNF113A complex was treated with or without lamda phosphatase, as indicated. E3 ligase assays were performed in the presence of the methylated β-globin pre-mRNA as in **(A)** (n=3 independent replicates).

These data suggested that RNF113A or an associated factor may recognize methyl adducts in RNA to regulate RNF113A E3 activity. We performed RNA-immunoprecipitation followed by mass spectrometry (RIP-MS) from cells expressing WT Flag-tagged RNF113A, in the presence or absence of MMS treatment. Upon MMS treatment, association between RNF113A and methylated RNA increased significantly (Figure 5C). Indeed, RNAs containing the three major MMS-induced modifications (m3C, m1A, and m7G) were associated with RNF113A during alkylation. We reasoned that this association between RNF113A and alkylated RNA would coincide with an increased binding between RNF113A and the active form of the spliceosome, which it is known to associate with in the active spliceosome (Zhang et al., 2018). By immunoprecipitation, we found markedly increased association between RNF113A and the spliceosomal protein BRR2 after MMS-induced alkylation (Figure 5D), which we previously showed is a potential substrate for RNF113A (Brickner et al., 2017).

During the course of analyzing RNF113A purified from human cells, we consistently noticed that RNF113A migrated slower than its calculated molecular weight (38 kDa untagged, 42 kDa with dual Flag and HA tags). Furthermore, its migration appeared further retarded post MMS treatment (see Figure 5D). These observations hinted that RNF113A is phosphorylated and its phosphorylation status is altered in response to alkylation stress. Indeed, phosphoproteomics analysis indicated that the protein is phosphorylated at multiple residues (see below). These findings were further confirmed through the use of the Phos-tag acrylamide reagent, which showed the mobility of the protein to be retarded relative to a phosphatase-treated counterpart (Supplemental Figure S6C). We note that these findings are in agreement with those of the accompanying manuscript, for which RNF113A is found to be a phosphoprotein regulated by the PP4 phosphatase (see Lukonovic et al.). However, while these observations establish RNF113A as a phosphoprotein, whether this post-translational modification plays a role during its response to alkylation damage is not immediately clear. Therefore, we tested whether phosphorylation of RNF113A is a prerequisite for its activation by alkylated RNA. Notably, pre-treatment of the purified RNF113A complex with lambda phosphatase markedly reduced its ability to respond to alkylated RNA, suggesting that its phosphorylation is important for its function in response to damaged RNA (Figure 5E).

### RNF113A phosphorylation is required for alkylation induced autoubiquitination and ASCC3 recruitment

To determine which sites are phosphorylated on RNF113A, we performed phosphoproteomics on the protein isolated from HeLa cells before and after MMS treatment. We found multiple sites throughout the protein that were potentially phosphorylated under both conditions (Supplemental Figure S7A). Data from the accompanying manuscript suggested that PP4 is repelled from RNF113A when the latter is methylated at K20 (see Lukonovic et al.); therefore, we focused on phosphorylation sites primarily within the N-terminal region of RNF113A. We mutated serine 6 to alanine (S6A), as well as four additional serine residues close to the N-terminus (S43A, S45A, S46A, and S47A, termed N4 mutant), at least two of which we found to be phosphorylated by MS (Figure 6A and Supplemental Figure S7A). In addition, we targeted two threonine residues within the RING domain (T292V and T293V) which were predicted phosphorylation sites *in silico* (http://gps.biocuckoo.cn). We then assessed MMS-induced autoubiquitination of these forms of RNF113A using the TUBE assay. Neither the S6A mutant alone, nor the N4 mutant significantly affected RNF113A autoubiquitination (Figure 6B). However, the combination of the S6A and N4 mutants (which we termed N5) markedly reduced RNF113A autoubiquitination in response to MMS, suggesting the functional significance of this modification in the damage response. Analysis of this mutant protein using a phos-tag gel suggested that these residues were at least partially phosphorylated even in the absence of exogenous alkylation, consistent with our MS data (Supplemental Figure S7B). Functionally, the RNF113A N5 mutant was still capable of associating with nuclear speckle bodies in a manner which mirrored the WT RNF113A (Figure S7C-S7D). The N5 mutant, as well as the S6A mutant also behaved as WT RNF113A for association with alkylated nucleotides as determined by RIP-MS, suggesting that the protein is otherwise functional and not misfolded (Figure 6C).

**Figure 6.**
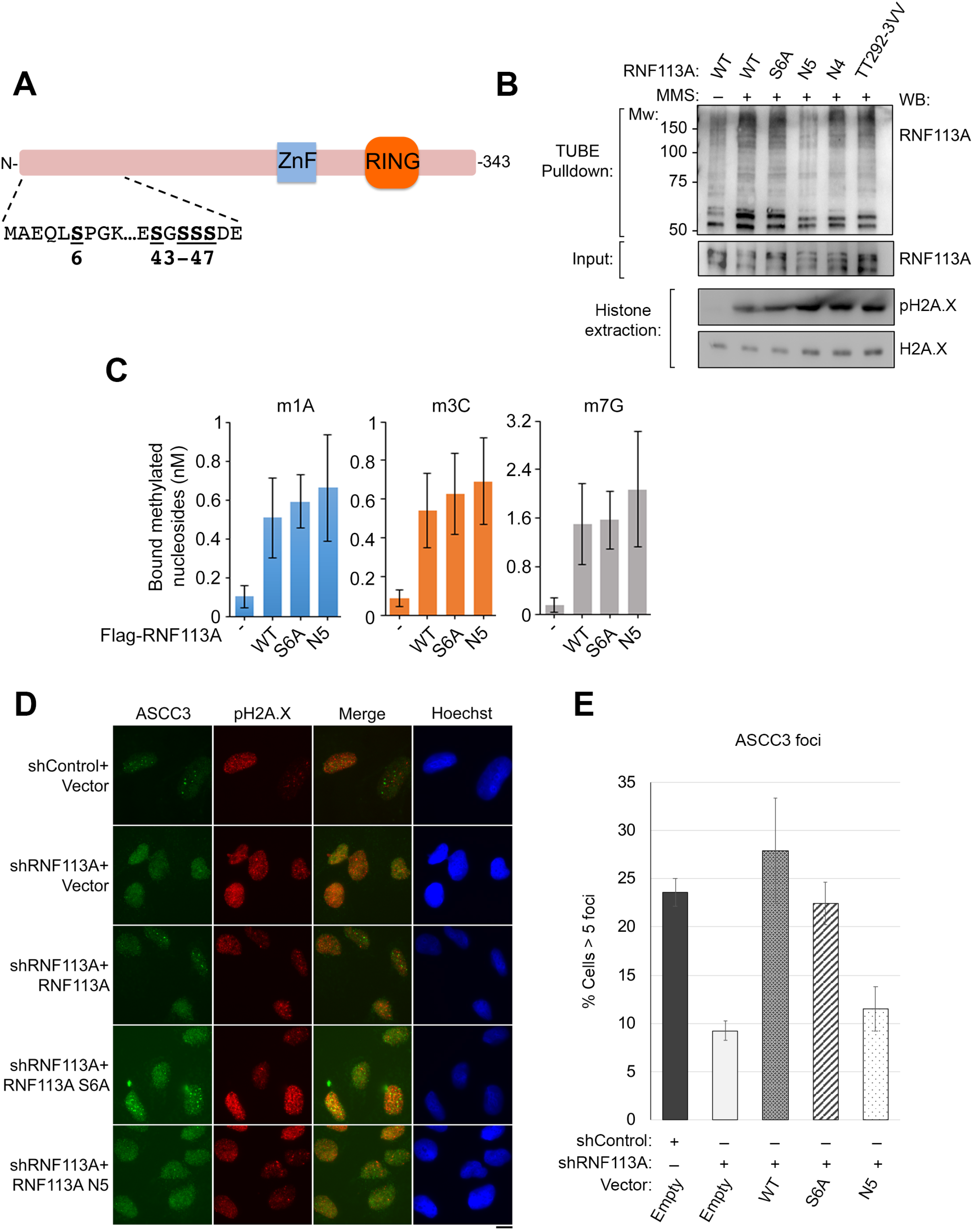
Phosphorylation of RNF113A is required for MMS-induced autoubiquitination and ASCC recruitment. **(A)** Schematic of RNF113A and its potential N-terminal phosphorylation sites targeted for mutagenesis. **(B)** HeLa-S cells expressing WT RNF113A or the indicated RNF113A mutants were treated with MMS as indicated. Cell lysates were subjected to TUBE pulldown to determine RNF113A autoubiquitination. (n=2 independent replicates). **(C)** RIP-MS was performed from MMS-treated HeLa-S cells expressing the indicated Flag vectors. Purified RNAs were digested and the methylated nucleosides were analyzed by mass spectrometry. The results are shown as the mean ± S.D. (n=3 independent replicates). **(D)** U2OS cells expressing the indicated shRNAs were rescued with the empty vector or the indicated Flag-RNF113A vectors. After MMS treatment, cells were processed for immunofluorescence staining using the indicated antibodies. Scale bar, 10 μm. **(F)** Quantitation of **(E)**. The results are shown as the mean ± S.D (n=2 replicates).

Our data suggested that RNF113A function may be governed by these N-terminal phosphorylation sites. To test the role of these sites in cells, we assessed MMS-induced ASCC3 foci formation in cells knocked down for endogenous RNF113A and reconstituted with WT RNF113A or specific phosphorylation mutants. While the WT and S6A mutant proteins were able to rescue ASCC3 foci formation, the N5 mutant was not capable of doing so (Figure 6E-6F and Supplemental Figure S7F). Altogether, this data strongly supports the model that RNF113A is a phosphorylated E3 ubiquitin ligase capable of responding to alkylated RNA in order to recruit the ASCC complex.

## Discussion

We had previously identified a ubiquitin-dependent signaling mechanism by which the ASCC-ALKBH3 repair complex is specifically recruited during alkylation damage to promote lesion reversal (Brickner et al., 2017). Recruitment of the helicase ASCC3 and the repair enzyme ALKBH3 is promoted by the accessory protein ASCC2, which recognizes RNF113A-mediated ubiquitination. Yet, the upstream events that result in complex recruitment, and how alkylation damage in particular activates this pathway, remains unanswered. In this study, we provide multiple lines of evidence that aberrant RNA methylation serves as a critical signal for the recruitment of this repair complex (Figures 1-2). Consequentially, the recruitment of ASCC appears to promote downregulation of damaged transcripts (Figure 3). We further show that alkylation damage selectively activates RNF113A and that alkylated RNA is sufficient to induce RNF113A E3 ligase activity (Figures 4-5). Finally, phosphorylation of RNF113A at specific residues is a prerequisite for activation of this E3 ligase and ASCC recruitment during alkylation damage (Figure 6).

Although relatively unexplored, recent work has begun to highlight the importance of RNA in propagating DNA repair, particularly for promoting homologous recombination during double-strand break repair (Bader et al., 2020; Mazina et al., 2017). While these studies have focused on the role of RNA-associated proteins, transcripts and R-loops in mediating repair, our study highlights a novel mechanism by which aberrant RNA methylation propagates damage response signaling to recruit the ASCC repair complex. Indeed, RNA alkylation is both necessary, as overexpression of an RNA-specific demethylase reduced ASCC2 recruitment in response to alkylation damage (Figure 1), and sufficient, as overexpression of an RNA methyltransferase that deposits a damage-associated methyl mark alone results in ASCC complex recruitment (Figure 2). While these “aberrant” methylations are strongly induced by alkylating agents, we should note that they are thought to exist in certain mRNAs, albeit in low amounts under normal cellular conditions, such as internal m7G sites within certain mRNAs (Zhang et al., 2019), which may in turn activate this pathway. Thus, it is plausible that even in unperturbed cells, this pathway is nominally responding to repair alkylated bases. Indeed, the basal phosphorylation of RNF113A in the absence of exogenous damage permits the pathway to be activated quickly in response to such endogenous events.

How cells shut down transcription, both locally and globally, in response to DNA damage is becoming an area of increasing interest. Here, we show that aberrant alkylation damage, either by alkylating agents such as MMS or overexpression of METTL8, results in repression of certain target genes (Figure 3). These alkylation-repressed transcripts are partially derepressed upon loss of either ASCC3 or ASCC2. These findings are consistent with previous results that suggest that the ASCC complex shuts down transcription globally upon UV damage (Williamson et al., 2017). We should note that while the majority of altered transcripts are repressed, transcription of RNA binding proteins and the unfolded protein response is upregulated, suggesting that these additional pathways may play a role in the cellular response to alkylation damage.

Consistent with our previous findings, we demonstrate that RNF113A E3 ligase activity is specifically stimulated by alkylation damage (Figure 4). Loss of RNF113A specifically sensitized cells to alkylation damage, but not to other DNA damaging agents. Thus, stimulation of RNF113A ligase activity functionally serves as a ‘rheostat’ for alkylation damage. This phenomenon appears to be specific to this alkylation repair pathway, as MMS did not dramatically induce the autoubiquitination of RNF8 or RNF168, two well-characterized ligases involved in double-strand break repair (Schwertman et al., 2016). It is notable that these latter E3 ligases appear to be regulated primarily by recruitment to sites of DNA double-stranded break-induced foci, whereas RNF113A is constitutively associated with nuclear speckle bodies (Brickner et al., 2017).

Our data indicates that post-translational modification of RNF113A by phosphorylation is needed to initiate this alkylation repair pathway. Phosphorylation of RNF113A appears to be required for its basal E3 ligase activity, as well as its ability to be further activated by alkylated RNA (Figures 5-6). Previous reports have identified a number of E3 ligases that can be activated by phosphorylation events, often through conformational changes (Durcan and Fon, 2015; Gallagher et al., 2006; Smith et al., 2009). Our findings are consistent with evidence in the accompanying manuscript which further bolsters phosphorylation-dependent mechanism of RNF113A regulation (see Lukonovic et al.). This latter work strongly suggests that the phosphatase PP4 acts on RNF113A to repress its E3 activity in a manner that is in turn regulated by SMYD3-induced methylation. Interestingly, our work indicates that a minimum of two distinct phosphorylation events are necessary for RNF113A activation: a phosphorylation event at S6 and another event among serine residues S43 and S45-S47. Consistent with this observation, S6 contains a canonical kinase consensus sequence for the CDK family of kinases, while the sequence context of S43 and S45-47 fits the consensus sequence for casein kinase 2 (CK2; Figure 6A). It is therefore likely that the coordination of multiple kinases, including a CDK, is necessary for the activation of RNF113A. Further investigation into the identity of the kinase(s) is required. Since loss of RNF113A phosphorylation prevented proper recruitment of the ASCC complex (Figure 6), the use of kinase inhibitors to prevent RNF113A activation represents an attractive option to increase alkylation sensitivity of cancer cells. The accompanying manuscript also suggests that SMYD3 inhibition may serve a similar function and improve alkylation damage responses (see Lukonovic et al.).

Our previous findings indicate that the ASCC-ALKBH3 complex serves as a repair complex (Dango et al., 2011; Brickner et al., 2017), whose loss results in impaired repair kinetics of alkylated lesions on DNA. As such, our work unveils a novel mechanism by which DNA repair complexes are initially recruited through RNA damage to sites of transcriptionally active, damaged regions. However, recent evidence strongly suggests that the ASCC complex promotes ribosome quality control in the cytosol, a pathway that is also activated upon alkylation damage during protein translation (Juszkiewicz et al., 2020; Matsuo et al., 2020). Therefore, our work has potentially unveiled the adaptation of RNA damage recognition as a novel mechanism to recruit this complex to damaged, transcriptionally active regions of the nuclear genome.

## Acknowledgments

We wish to thank Yang Shi, Alessandro Vindigni, Hani Zaher, and members of the Structural Cell Biology of DNA Repair Machines (SBDR) program project for their advice on this manuscript. V.L. is a recipient of the FRM postdoctoral fellowship (Fondation pour la Recherche Médicale SPF201809006930). We acknowledge the Extreme Science and Engineering Discovery Environment (XSEDE), which is supported by NSF grant ACI-1548562. J.A.T. acknowledges support as a CPRIT Scholar in Cancer Research and Robert A. Welch Distinguished Chair in Chemistry. This work was supported by the ANR (ANR-16-CE11-0018 to N.R.), the INCa (PLBIO19-021 to N.R. and N.M.), the NIH (P01 CA092584 to J.A.T. and N.M., R01 CA193318 and R01 CA227001 to N.M.), the American Cancer Society (RSG-18-156-01-DMC to N.M.), and the Siteman Investment Program (to N.M.), and the Centene Personalized Medicine Initiative (to N.M.). C.H. is an investigator of the Howard Hughes Medical Institute.

## Author Contributions

J.R.B., N.T., R.R., C.O., V.L., J.M.S., A.G.C., A.G., and N.M. carried out cellular and biochemical experiments. A.B. performed bioinformatic analysis. L.Z. performed methylated lesion localization and analysis. C.H. supervised L.Z. J.A.T. supervised A.B. N.R. supervised V.L. and A.G.C. N.M. supervised the project and wrote the manuscript with J.R.B. and N.T., with input from all other authors.

## STAR Methods

### KEY RESOURCES TABLE

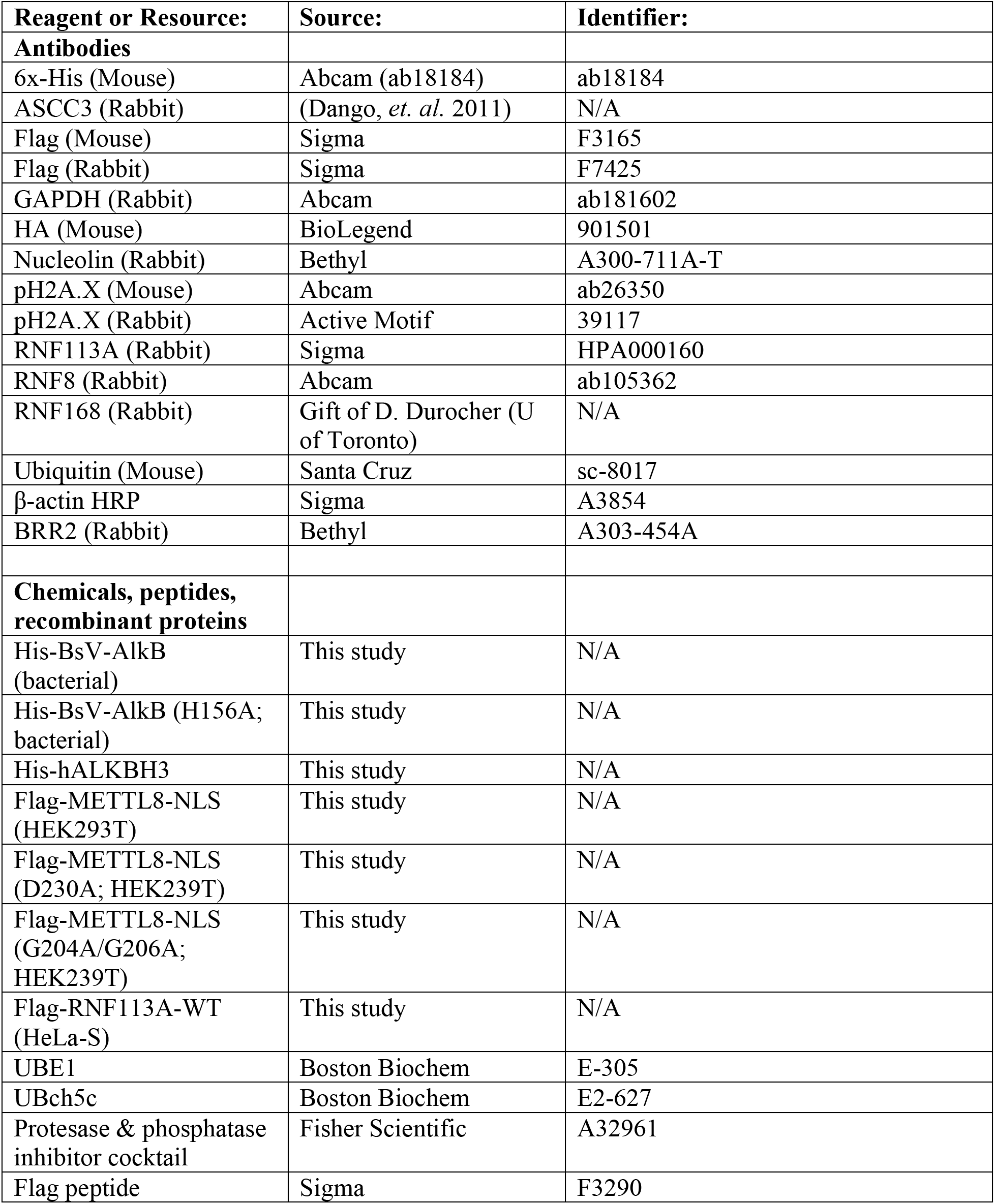
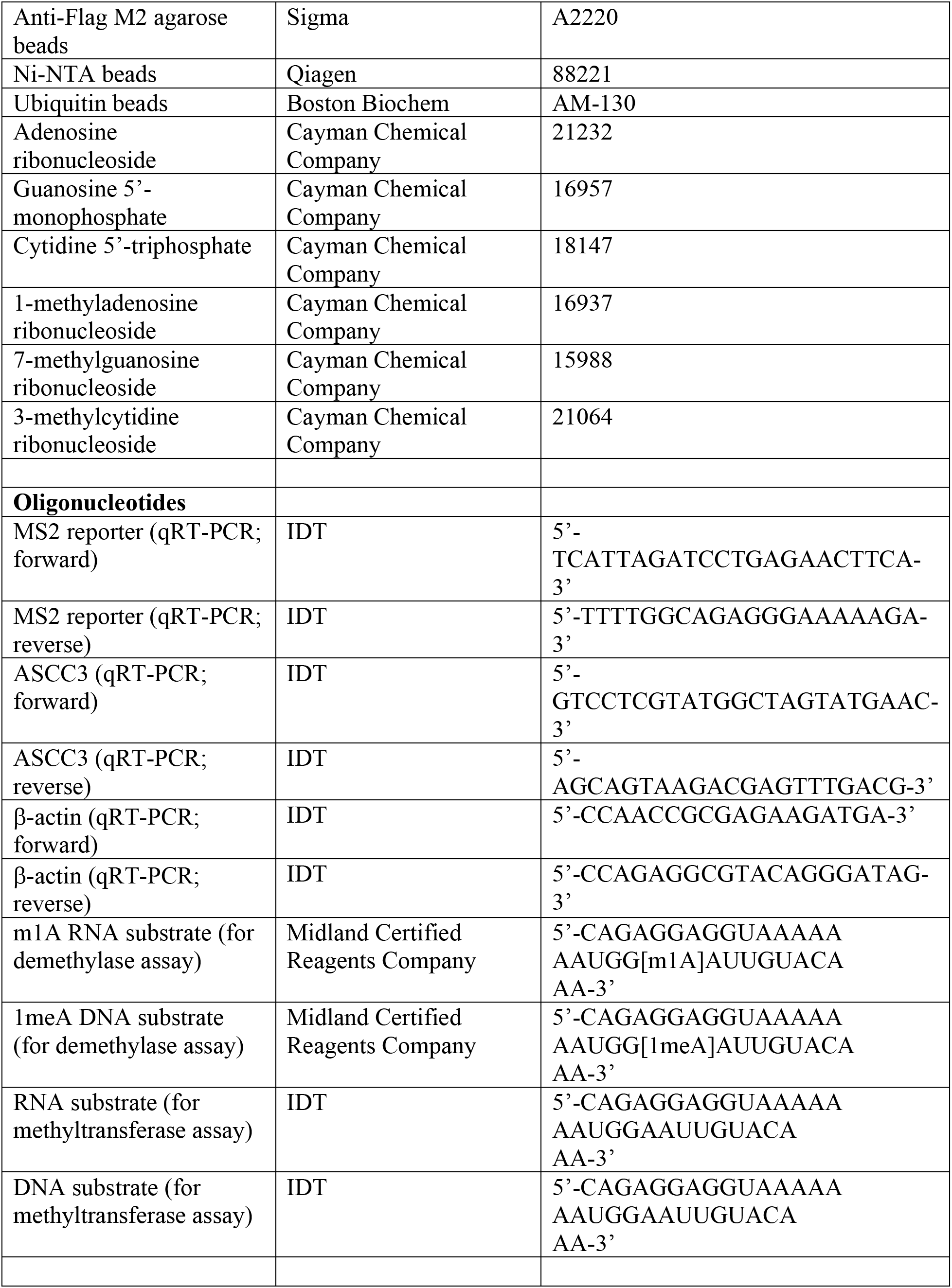
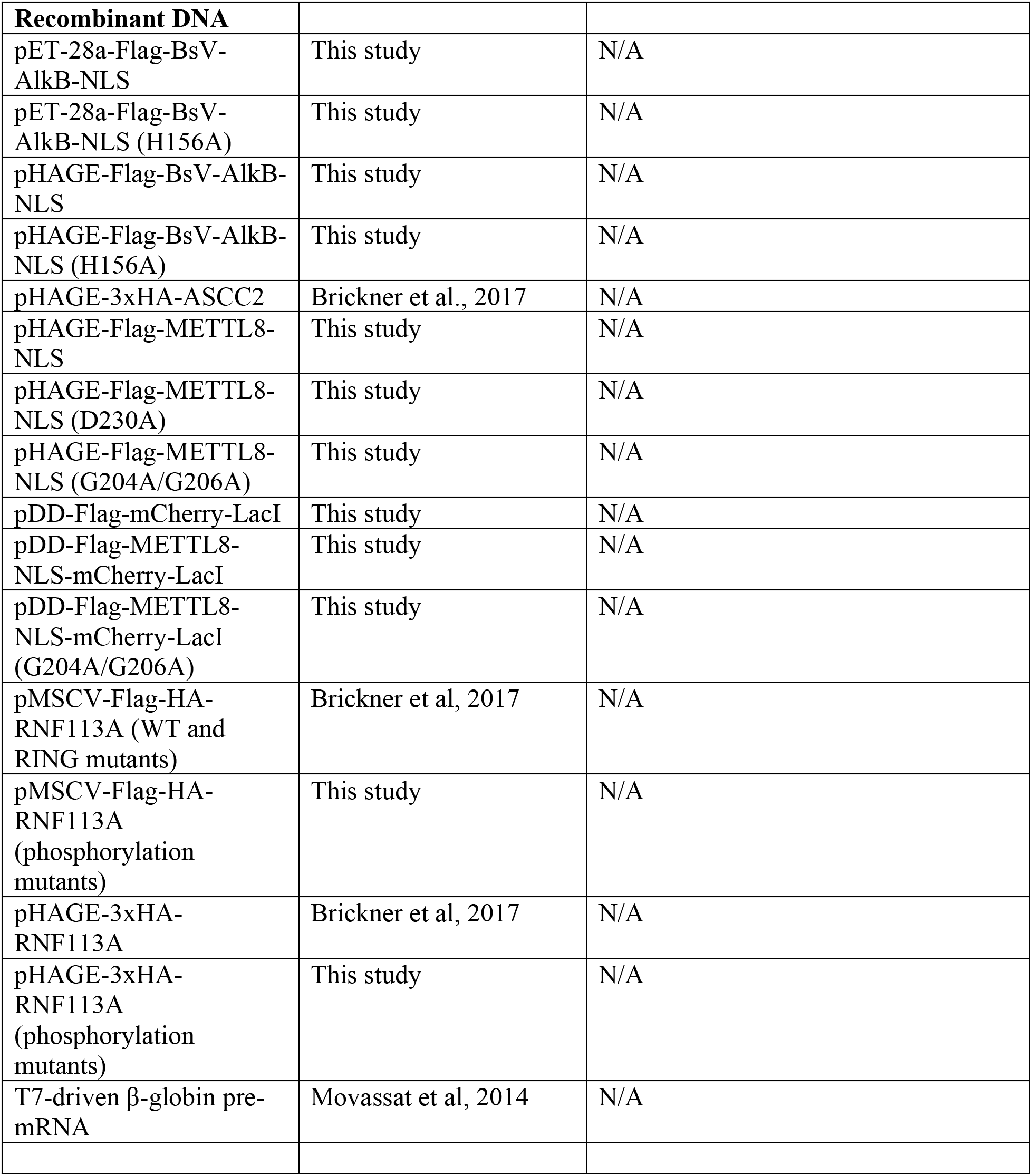

### CONTACT FOR REAGENT AND RESOURCE SHARING

Further information and requests for resources and reagents should be directed to and will be fulfilled by the Lead Contact, Nima Mosammaparast.

### EXPERIMENTAL MODEL AND SUBJECT DETAILS

#### Cell culture

All human cell lines were cultured in Dulbecco’s modified eagle medium (Invitrogen), supplemented with 10% fetal bovine serum (Atlanta Biologicals), 100 U/ml of penicillin-streptomycin (Gibco) at 37°C and 5% CO_2_.

### METHOD DETAILS

#### Plasmids

Plasmids containing human ASCC2 and RNF113A cDNAs were previously described (Dango et al., 2011). The BsV-AlkB cDNA(van den Born et al., 2008) was synthesized as a gBlock (IDT) after optimization of the sequence for human cell expression, fused to the SV40 NLS, and cloned into pENTR3-C. METTL8 was amplified from human cDNA, fused to the SV40 NLS, and cloned into pENTR-3C. For mammalian cell expression, cDNAs were subcloned into pHAGE-CMV-3xHA, pHAGE-CMV-Flag, pMSCV (no tag), or pMSCV-Flag-HA as needed by Gateway recombination(Brickner et al., 2017). For degron tagged expression, the mCherry-LacI sequence (from Addgene #18985) was subcloned into pLVX-PTuner (Takara Bio). To create the DD-Flag-METTL8-NLS-mCherry-LacI, Flag-METTL8-NLS was subcloned into the pLVS-PTuner-mCherry-LacI. For recombinant protein expression, cDNAs were subcloned into pET28a-Flag. Mutations were created by PCR-mediated mutagenesis or synthesized as gBlocks (IDT). For shRNA-mediated knockdowns, TRC pLKO.1 vectors were used as previously described (Brickner et al., 2017). For CRISPR/Cas9 mediated knockout of RNF113A, gRNA sequences were cloned into pLentiCRISPR-V2 (Addgene#52961). The T7 β-globin minigene construct for *in vitro* transcription of β-globin pre-mRNA was kindly provided by Dr. Klemens Hertel (University of California, Irvine), and was described previously(Movassat et al., 2014). All constructs derived by PCR or from gBlocks were confirmed by Sanger sequencing.

#### Cell culture and cell survival assays

293T, HeLa, HeLa-S, and U2OS cells (originally from ATCC) were cultured and maintained as previously described(Brickner et al., 2017). Cells were tested for mycoplasma at the Washington University Genome Engineering and iPSC Center. The U2OS ASCC3 knockout cell line created by CRISPR/Cas9 was previously described(Brickner et al., 2017). The U2OS 2-6-3 and 2-6-5 reporter cell lines(Janicki et al., 2004; Shanbhag et al., 2010) were a kind gift of Roger Greenberg (University of Pennsylvania). Preparation of viruses, transfection, and viral transduction were performed as described previously(Brickner et al., 2017). Knockdown experiments (using shRNA) and knockout experiments (using CRISPR/Cas9) were performed by infecting cells with the indicated lentivirus and selecting with puromycin (1 μg/ml) for 48-72 hours. For ASCC foci rescue experiments, cells were transduced with the indicated pMSCV retroviral vector. For DNA damaging agent survival assays using HeLa cells, 2,000 cells/well were cultured overnight in 96-well plates in 100 μl media. Cells were then exposed to medium containing the indicated concentration of methyl methanesulphonate (MMS; Sigma) for 24 hours at 37 °C. The media was then replaced with normal media, and cell viability was assessed using the MTS assay (Promega) 72 hours after initial damaging agent exposure. For experiments involving camptothecin (CPT; Sigma), cells were exposed to medium containing the indicated concentration of the damaging agent in culture medium for 72 hours at 37 °C. Viability was then processed by MTS assay as above. All MTS-based survival experiments were carried out in technical quintuplicate.

#### Protein purification

Recombinant, His-tagged BsV-AlkB and ALKBH3 proteins were purified from Rosetta (DE3) cells using an ÄKTA-pure FPLC (GE Healthcare). Cells were resuspended in His-lysis buffer (50 mM Tris-HCl pH 7.3, 250 mM NaCl, 0.05% Triton X-100, 3 mM β-ME, 30 mM imidazole, and protease inhibitors) and lysed by sonication. After centrifugation and filtration, the extract was loaded onto a HisTrap HP column using a 50 ml Superloop (GE Healthcare). After extensive washing with lysis buffer, the protein was eluted using lysis buffer containing 400 mM imidazole. All recombinant proteins were dialyzed into TAP buffer. Flag-tagged METTL8 was purified from transiently transfected 293T cells by resuspension in Flag-lysis buffer (50 mM Tris-HCl pH 7.9, 150 mM NaCl, 10% glycerol 1.0% Triton X-100, 1 mM DTT, and protease inhibitors) and lysed by sonication. After incubation with Flag resin, beads were washed three times with 10 ml of Flag-lysis buffer and twice with 4 mL Flag-lysis buffer. The protein was eluted with TAP wash buffer (50 mM Tris, pH 7.9, 100 mM KCl, 5 mM MgCl_2_, 0.2 mM EDTA, 0.1% NP-0, 10 % glycerol, 2 mM DTT) containing 0.4 mg/ml Flag peptide. Flag-tagged RNF113A was purified from HeLa-S cell nuclear extract as previously described (Brickner et al., 2017).

#### Alkylated RNA preparation

*In vitro* transcription of β-globin pre-mRNA was performed by using HiScribe T7 *In Vitro* Transcription Kit (New England Biolabs). The *in vitro* transcribed pre-mRNA was treated with dimethyl sulfate (DMS) as described(Tijerina et al., 2007). 2 μM of the RNA was added into a 25 μl reaction mixture containing 0.65% DMS, 50 mM Na-MOPS pH 7.0, 10 mM MgCl_2,_ and incubate at room temperature for 10 minutes. Reactions were quenched with 475 μl quench solution (0.3 M sodium acetate, 30% β-Mercaptoethanol) prior to ethanol precipitation. RNA pellets were dissolved in water and concentration was determined by Nanodrop. The increase in methylation of the ribonucleosides was analyzed by quantitative LC-MS/MS, as described below.

#### Nucleic acid demethylase and methyltransferase assays

For demethylation assays, methylated oligonucleotides (RNA and DNA sequences shown in Key Resources Table) were purchased from Midland Certified Reagents Company (Midland, TX). Demethylation reactions were carried out using 60 pmol of substrate oligonucleotide in the presence of 20 pmol BsV-AlkB protein for 1 hour at 37°C in a 50 μl reaction mixture containing 50 mM HEPES–KOH pH 7.5, 2 mM ascorbic acid, 100 μM 2-oxoglutarate, 40 μM FeSO4 and 1 uL RNaseOUT recombinant ribonuclease inhibitor (ThermoFisher). Methyltransferase assays were carried out using 100 pmol of substrate oligonucleotide (IDT; see Key Resources Table for sequence) in the presence of ~0.5μg of immunopurified Flag-METTL8 for 2 hours at 37°C in a 40 μl reaction mixture containing 50 nM HEPES pH 7.5, 3 mM MgCl_2_, 0.5 mM S-adenosylmethonine and 1 μL RNaseOUT recombinant RNase inhibitor. The products for both types of reactions were digested to nucleosides and analyzed by quantitative LC-MS/MS, as described below.

#### RNA immunoprecipitation (RIP)

The procedure was followed as previously described (Wang et al., 2014), with minor modifications. HeLa-S cells expressing empty pMSCV vector, Flag-RNF113A (WT), or Flag-RNF113A ZnF point mutant (K218A/K219A) grown in 15-cm dishes were treated with or without 5 mM MMS for 1 h. Cells were harvested and the pellets were lysed with 2 volumes of lysis buffer (10 mM HEPES pH 7.5, 150 mM KCl, 0.5% NP-40, 2mM EDTA, 0.5mM Zn(NO_3_)_2_, 10 mM β-ME, protease inhibitors and 400 U/ml RNase inhibitor), incubated on ice for 5 minutes then shock-frozen at −80°C for 30 minutes. Cell lysates were thawed on ice and centrifuged at 15,000g for 15 minutes. The supernatant was immunoprecipitated with M2 agarose beads at 4°C for 4hrs. Beads were washed eight times with 1 ml ice-cold NT2 buffer (50 mM HEPES pH 7.5, 200 mM NaCl, 0.05% NP-40, 2mM EDTA, 0.5mM Zn(NO_3_)_2_, 10 mM β-ME and 200 U/ml RNase inhibitor) and eluted with 5 packed bead volumes of NT2 buffer containing 0.4 mg/ml Flag peptide. The supernatant was mixed with 1 ml TRIzol for RNA extraction according to the manufacture’s protocol, then digested for nucleoside mass spectrometry analysis.

#### Nucleoside Mass Spectrometry

Samples were digested to nucleosides at 37°C overnight with Nuclease S1 from *Aspergillus oryzae* (Sigma-Aldrich), followed by dephosphorylation with bovine alkaline phosphatase (Sigma Aldrich) at 37°C for 1 hour. Chromatographic separation was performed using an Agilent 1290 Infinity II UHPLC system with a ZORBAX RRHD Eclipse Plus C18 2.1 x 50 mm (1.8 um) column. The mobile phase consisted of water and methanol (with 0.1% formic acid) run at 0.5 ml/min. For DNA nucleosides, the run started with a 3 min gradient of 2-8% methanol, followed by a 2 min gradient of 8-98% methanol, 98% methanol was maintained for 4 min, followed by re-equillibration to 2% methanol over 1 min. For RNA nucleosides, the run started with a 3 min gradient of 2-8% methanol, followed by a sharp increase to 98% methanol which was maintained for 4 min. Mass spectrometric detection was performed using an Agilent 6470 Triple Quadrupole system operation in positive electrospray ionization mode, monitoring the mass transitions 266.13/150 (for m1dA), 252.11/136 (for dA), 282/150 (for m1A), 268.1/136 (for A), 242.1/126.1 and 242.1/109 (for m3dC), 228.1/112 (for dC), 258/126 (for m3C), 244.1/112 (for C), 298/166 (for m7G), and 284.2/152 (for G).

#### Phosphorylation site identification by mass spectrometry

The excised gel band containing Flag-RNF113A was cut into approximately 1 mm pieces. Gel pieces were then subjected to a modified in-gel trypsin digestion procedure (Shevchenko et al., 1996). Gel pieces were washed and dehydrated with acetonitrile for 10 minutes. Acetonitrile was removed and gel pieces were completely dried in a speed-vac. The gel was rehydrated with 50 mM ammonium bicarbonate solution containing 12.5 ng/μl modified sequencing-grade trypsin (Promega) at 4°C. After 45 minutes, the excess trypsin solution was removed and replaced with 50 mM ammonium bicarbonate solution to just cover the gel pieces. Samples were incubated at 37°C overnight. Peptides were extracted by removing the ammonium bicarbonate solution, followed by one wash with a solution containing 50% acetonitrile and 1% formic acid. The extracts were dried in a speed-vac. For LC-MS/MS, peptides were resuspended in 6μl 1% formic acid and analyzed on an Orbitrap Fusion mass spectrometer (Thermo Fisher Scientific) equipped with a Proxeon Easy nLC 1000 for online sample handling and peptide separations. A portion of the peptides was loaded onto a 100 μm inner diameter fused-silica micro capillary with a needle tip pulled to an internal diameter less than 5 μm. The column was packed in-house to a length of 35 cm with a C18 reverse phase resin (GP118 resin 1.8 μm, 120 Å, Sepax Technologies). The peptides were separated using a 120 minute linear gradient from 3% to 25% buffer B (100% ACN + 0.125% formic acid) equilibrated with buffer A (3% ACN + 0.125% formic acid) at a flow rate of 600 nL/min across the column. The scan sequence for the Fusion Orbitrap began with an MS1 spectrum (Orbitrap analysis, resolution 120,000, 400−1400 m/z scan range, AGC target 2 × 10^5^, maximum injection time 100 ms, dynamic exclusion of 30 seconds). The most intense precursor from each MS1 scan was selected for MS2 analysis. Peptides were isolated in the quadrupole using an isolation window of 0.5 Da, fragmented by HCD with a collision energy of 35%, and analyzed in the orbitrap (resolution 60,000, 350-1400 m/z scan range, AGC target 2 × 10^4^) with a maximum injection time of 150 ms. The remaining peptides were analyzed over an 85 minute linear gradient using a “most intense” method in which MS1 collection was the same as above, but MS2 analysis after quadrapole isolation (isolation window 0.7 Da) was performed using CID activation with a collision energy of 30% followed by fragment ion detection in the ion trap (350-1400 m/z scan range, 35 ms maximum injection time, AGC target 1 × 10^4^). Data analysis was performed using a suite of in-house software tools for .RAW file processing and controlling peptide and protein level false discovery rates, assembling proteins from peptides, and phosphopeptide identification (Huttlin et al., 2010).

#### RNA Sequencing and Analysis

RNA was purified using the Qiagen RNeasy mini kit. Samples were prepared according to library kit manufacturer’s protocol, indexed, pooled, and sequenced on an Illumina HiSeq. Basecalls and demultiplexing were performed with Illumina’s bcl2fastq software and a custom python demultiplexing program with a maximum of one mismatch in the indexing read. RNA-seq reads were then aligned to the Ensembl release 76 top-level assembly with STAR version 2.0.4b(Dobin et al., 2013). Gene counts were derived from the number of uniquely aligned unambiguous reads by Subread:featureCount version 1.4.5(Liao et al., 2014). Isoform expression of known Ensembl transcripts were estimated with Salmon version 0.8.2(Patro et al., 2017). Sequencing performance was assessed for the total number of aligned reads, total number of uniquely aligned reads, and features detected. The ribosomal fraction, known junction saturation, and read distribution over known gene models were quantified with RSeQC version 2.3 (Wang et al., 2012).

All gene counts were then imported into the R/Bioconductor package EdgeR(Robinson et al., 2010) and TMM normalization size factors were calculated to adjust for samples for differences in library size. Ribosomal genes and genes not expressed in the smallest group size minus one samples greater than one count-per-million were excluded from further analysis. The TMM size factors and the matrix of counts were then imported into the R/Bioconductor package Limma(Ritchie et al., 2015). Weighted likelihoods based on the observed mean-variance relationship of every gene and sample were then calculated for all samples with the voomWithQualityWeights(Liu et al., 2015). The performance of all genes was assessed with plots of the residual standard deviation of every gene to their average log-count with a robustly fitted trend line of the residuals. Differential expression analysis was then performed to analyze for differences between conditions and the results were filtered for only those genes with Benjamini-Hochberg false-discovery rate adjusted *p*-values less than or equal to 0.05. For each contrast extracted with Limma, global perturbations in known Gene Ontology (GO) terms and KEGG pathways were detected using the R/Bioconductor package GAGE(Luo et al., 2009) to test for changes in expression of the reported log 2 fold-changes reported by Limma in each term versus the background log 2 fold-changes of all genes found outside the respective term. Heatmaps were generated using the Heatmapper online tool (heatmapper.ca). All RNA-Seq primary data and metadata is to be deposited in the Gene Expression Omnibus (https://www.ncbi.nlm.nih.gov/geo) and is available upon request.

#### Immunofluorescence microscopy

All immunofluorescence microscopy was done as previously described (Brickner et al., 2017), with minor modifications. U2OS cells expressing NLS-tagged METTL8 fusions were washed with 1× PBS prior to fixation with 3.2% paraformaldehyde. For YFP-MS2 analysis, the U2OS 2-6-3 expressing degron-tagged LacI fusion proteins were incubated with 300 nM Shield1 ligand (Takara Bio) and 1 μg/ml doxycycline for 24 hours, then washed with 1× PBS prior to fixation as above. For ASCC localization, U2OS 2-6-5 cells were treated as the 2-6-3 cells but were extracted with 1× PBS containing 0.2% Triton X-100 and protease inhibitor cocktail (Pierce) for 10-20 minutes on ice prior to fixation. For MMS-induced foci analysis, U2OS cells were treated with 500 μM MMS in complete medium at 37°C for six hours, washed with 1× PBS, then extracted and fixed as above. All cells were then washed extensively with IF Wash Buffer (1× PBS, 0.5% NP-40, and 0.02% NaN_3_), then blocked with IF Blocking Buffer (IF Wash Buffer plus 10% FBS) for at least 30 minutes. Primary antibodies were diluted in IF Blocking Buffer overnight at 4°C. After staining with secondary antibodies (conjugated with Alexa Fluor 488 or 594; Millipore) and Hoechst 33342 (Sigma-Aldrich), where indicated, samples were mounted using Prolong Gold mounting medium (Invitrogen). Epifluorescence microscopy was performed on an Olympus fluorescence microscope (BX-53) using an ApoN 60X/1.49 NA oil immersion lens or an UPlanS-Apo 100X/1.4 oil immersion lens and cellSens Dimension software. Raw images were exported into Adobe Photoshop, and for any adjustments in image contrast or brightness, the levels function was applied. For foci quantitation, at least 100 cells were analyzed in triplicate, unless otherwise indicated.

#### His-ubiquitin pulldown assays

His-tagged ubiquitin was immunoprecipitated after denaturation as described previously(Gajjar et al., 2012; Xirodimas et al., 2001), with minor modifications. Briefly, HeLa cells expressing Flag-HA-RNF113A WT, Flag-HA-RNF113A I264A or Flag-HA-RNF113A ΔRING were transfected with His-Ub. At ~42 hours after transfection, cells were exposed to various DNA damaging agents for an additional 6 hours (500 μM MMS, 10mM HU, 20 μM bleomycin, 1 μM CPT). Cells were washed twice with cold 1x PBS and lysed in 1ml of Lysis buffer (6 M guanidinium-HCl, 100 mM Na_2_HPO_4_/NaH_2_PO_4_, 10 mM Tris-HCl pH 8.0, 5 mM imidazole, 10 mM β-mercaptoethanol, and protease inhibitors). Cells were sonicated on ice for 10 seconds twice and centrifuged at 11K rpm at 4°C for 10min. The supernatant was collected in a new tube and 4ml of Lysis buffer was added. Ni-NTA-agarose beads were washed four times with Lysis buffer, added to the lysate and incubated for 4 hours at room temperature with rotation. Samples were washed for 5min at room temperature once with Lysis buffer, once with Wash buffer (8 M urea, 100 mM Na_2_HPO_4_/NaH_2_PO_4_, 10 mM Tris-HCl pH 6.8, 5 mM imidazole, 10 mM β-mercaptoethanol, and protease inhibitors), and twice with Wash Buffer +0.1% Triton X-100. Beads were then incubated with Elution Buffer (50 mM Tris-HCl pH 7.3, 250 mM NaCl, 400mM imidazole, 0.05% Triton X-100, 3 mM β-mercaptoethanol, and protease inhibitors) overnight at 4°C. After elution, 4x Laemmli Buffer was added, and samples were analyzed by Western Blotting.

#### Tandem ubiquitin binding element (TUBE) assays

HeLa-S cells (~6-8 × 10^7^) grown in a spinner flask were treated with the indicated genotoxic agents (1mM MMS, 10mM hydroxyurea, 20 μM bleomycin, 1 μM camptothecin, 200 μM H_2_O_2_) for four hours at 37°C. The cells were collected by centrifugation, and washed with ice-cold PBS and frozen at −80°C as two cell pellets. Each cell pellet was resuspended in 10ml TUBE lysis buffer (50 mM Tris·HCl, pH 7.5, 1 mM EGTA, 1 mM EDTA, 1% (v/v) Triton X-100, and 0.27 M sucrose) containing freshly added 100mM iodoacetamide, protease and phosphatase inhibitors, then rotated at 4°C for one hour for lysis. The extract was spun at 6500 rpm for 30 minutes. Supernatant was transferred to a fresh tube and spun at 6500 rpm for 5 minutes. A fraction of the spun extract was kept as input, and the rest was rotated overnight at 4°C with 50μl commercial ubiquilin-conjugated TUBE beads (Boston Biochem) or HALO-conjugated TUBE beads (Zhao et al., 2018). The beads were washed three times with 10 ml high salt TAP buffer (50 mM Tris-HCl, pH 7.9, 300 mM KCl, 5 mM MgCl2, 0.2 mM EDTA, 0.1% NP-0, 10% glycerol, 2 mM mercaptoethanol, 0.2 mM PMSF) and once with 1 mL low salt TAP buffer (50 mM Tris-HCl, pH 7.9, 0 mM KCl, 5 mM MgCl2, 0.2 mM EDTA, 0.1% NP-0, 10% glycerol, 2 mM mercaptoethanol, 0.2 mM PMSF). Beads were resuspended in 50 μL Laemmli buffer and analyzed by Western blotting.

#### Ubiquitin ligase assays

Reactions analyzing ubiquitin chain polymerization were performed in ubiquitin ligase buffer (25 mM Tris pH 7.3, 25 mM NaCl, 10 mM MgCl_2_, 100 nM ZnCl_2_, 1 mM mercaptoethanol) containing 2 mM ATP and 10 μM of ubiquitin in a total volume of 20 μl. E1 activating enzyme (UBE1; Boston Biochem) was used at 31.25 nM, and E2 ubiquitin conjugating enzymes (Ubch5c; Boston Biochem) were used at 0.625 μM. Flag-tagged-RNF113A purified from HeLa-S cells with or without treatment with MMS was added to each reaction and incubated at 37°C for 3 hours. For the reactions incubated with nucleic acid, Flag-tagged-RNF113A or RNF113A mutant proteins purified from HeLa-S cells was used at 0.1 μM. *In vitro*-transcribed β-globin pre-mRNA with or without treatment with DMS, as well as DNA or RNA oligonucleotides with or without single m1A in the sequences were added as indicated concentrations and incubated at 37°C for 1.5 hours. Reactions were stopped with 20 μl of Laemmli buffer, analyzed by SDS-PAGE, and Western blotted.

#### Statistical Analyses

All *p*-values related, except those related to RNA-Seq box plots, were calculated by unpaired, two-tailed Student’s *t*-test. RNA-Seq box plot *p*-values were determined by an exact permutation of the Wilcoxon-Mann-Whitney as described(Marx et al., 2016). All error bars represent the standard deviation of the mean, unless otherwise noted.

## Supplemental Figure Legends

**Supplemental Figure S1. Characterization of the Blueberry Scorch Virus (BsV) AlkB demethylase. (A)** Wildtype and a catalytic mutant (H156A) form of His_6_-tagged BsV-AlkB protein purified from bacteria was separated on 4%-12% SDS-PAGE gel and stained with Coomassie brilliant blue (CBB). Positions of molecular weight markers (Mw) are shown on the left. **(B)** An RNA oligonucleotide containing a single 1-methyladenine was subjected to demethylation using the BsV-AlkB proteins from **(A)**. Sample ion intensity chromatograms from LC-MS/MS analysis of the digested nucleoside products is shown. **(C)** Demethylation assays were performed as in **(B)**, except that a DNA oligonucleotide with the identical sequence was used as a substrate for the reaction. **(D)** Immunofluorescence analysis of U2OS cells expressing Flag-vector, Flag-BsV-AlkB-NLS (WT), or the Flag-BsV-AlkB-NLS (H156A). Cells were stained for the Flag antigen and the nuclei were counterstained with Hoechst. Scale bar, 10μm.

**Supplemental Figure S2. Characterization of the METTL8 RNA methyltransferase. (A)** Schematic of the overall structure of the human METTL8 methyltransferase (top). Alignment of the predicted S-adenosylmethionine (SAM) binding domain is shown (bottom), with arrows indicating the conserved residues (G204//206 and D230) chosen for mutagenesis. **(B)** Immunofluorescence analysis of U2OS cells expressing Flag-METTL8-NLS (WT), Flag-METTL8-NLS (D230A), or Flag-METTL8-NLS (G204/206A). Cells were stained for the Flag antigen, nucleolin, and the nuclei were counterstained with Hoechst. Arrows indicate nucleoli. Scale bar, 10μm. **(C)** Flag-GFP, Flag-METTL8-NLS (WT), and Flag-METTL8-NLS (G204/206A) were purified from 293T cells, separated on 4%-12% SDS-PAGE gel and silver stained. **(D)** Total nuclear intensity of the Flag antigen was quantified from cells expressing the indicated vectors (corresponding to experiments in Figure 2C-2D). N = 3 replicates and error bars indicate ±S.D. of the mean.

**Supplemental Figure S3. Transcriptome analysis during alkylation damage. (A)** and **(B)** Top 25 altered GO cellular components analysis **(A)** and GO molecular functions analysis **(B)** comparing RNA-Seq transcriptome data from MMS-treated versus untreated cells (n = 3 biological replicates per condition). **(C)** and **(D)** Volcano plot of RNA-Seq transcriptome analysis comparing WT U2OS with ASCC3 KO cells **(C)** or WT U2OS with ASCC2 KO cells **(D)** in the absence of MMS treatment. Genes upregulated or downregulated more than ±2 log_2_ fold change (FC) in the knockout condition are highlighted in green and red, respectively. **(E)** Heatmap of MMS downregulated genes in WT versus ASCC2 KO cells.

**Supplemental Figure S4. Aberrant RNA methylation mediates transcriptional downregulation. (A)** Relative quantitative RT-PCR analysis of the MS2 transcript was performed in the U2OS reporter system transduced with the DD-Flag-METTL8-mCherry-LacI vector under the indicated conditions. N = 3 replicates and error bars indicate ±S.D. of the mean. **(B)** His_6_-tagged human ALKBH3 protein purified from bacteria was separated on 4%-12% SDS-PAGE gel and stained with Coomassie brilliant blue (CBB) is shown on the left. Quantitative RT-PCR analysis (right) was performed with or without pre-incubation of the total RNA with ALKBH3 to demethylate m3C prior to qRT-PCR. N = 3 replicates and error bars indicate ±S.D. of the mean. **(C)** Quantitative RT-PCR analysis of ASCC3 was performed after infection with the indicated shRNAs. N = 3 replicates and error bars indicate ±S.D. of the mean. **(D)** Volcano plot of RNA-Seq transcriptome analysis comparing U2OS cells with or without induction of degron-tagged METTL8-NLS with Shield1 for 24 hours. Genes upregulated or downregulated more than ±1 log_2_ fold change (FC) in the +Shield1 condition are highlighted in green and red, respectively.

**Supplemental Figure S5. Selective activation of RNF113A by alkylation damage. (A)** HeLa cells expressing His-Ub and the indicated HA vectors were treated with MMS as shown. Whole cell extracts were then used for His-pulldowns using Ni-NTA beads under denaturing conditions. Lysate inputs as well as the pulldown material were analyzed by Western blot using the antibodies as shown. **(B)** TUBE assays were performed from HeLa-S cell extracts with the indicated genotoxic agents, as indicated. **(C)** Whole cell lysates from HeLa-S cells treated with the genotoxic agents were blotted with the pH2A.X and β-actin antibodies. **(D)** TUBE assays were performed from HeLa-S cell extracts with or without MMS treatment using general Ub TUBE or a K63-specific TUBE, as shown. Input lysates are shown on the left. **(E)** TAP-tagged RNF113A was affinity purified using Flag resin from HeLa-S cell nuclear extract with or without prior MMS treatment. The elute was separated on 4%-12% SDS-PAGE gel and silver stained. **(F)** E3 ubiquitin ligase assay was performed using the material from **(E)** and blotted with anti-ubiquitin or anti-Flag antibodies. **(G)** HeLa cells were transduced with the indicated gRNA vectors. Whole cell lysates were subsequently analyzed by Western blot using the antibodies as shown. **(H)** and **(I)** gRNA knockout of RNF113A was used for clonogenic survival assay using MMS **(H)** or camptothecin **(I)**.

**Supplemental Figure S6. Characterization of DMS treated pre-mRNA and RNF113A. (A)** *In vitro*-transcribed β-globin pre-mRNA was treated with 0.65% dimethyl sulfate (DMS) for 10 minutes. RNAs were purified and digested for the analysis of methylated nucleosides by LC-MS/MS. **(B)** E3 ubiquitin ligase assays were performed using RNF113A complex, in the presence or absence of the methylated or unmethylated β-globin pre-mRNA, as indicated. **(C)** Flag-RNF113A purified from HeLa-S cells was treated with calf intestinal phosphatase, as shown. The material was then analyzed on a Phos-tag gel and Western blotted using anti-Flag antibodies.

**Supplemental Figure S7. Characterization of RNF113A phosphorylation sites. (A)** Peptides identified and phosphorylation sites in RNF113A identified by LC-MS/MS. **(B)** Flag-RNF113A (WT, S6A, and N5 mutants) purified from HeLa-S cells were analyzed by Phos-tag gel and Western blotted as shown. **(C)** U2OS cells were transduced with the indicated HA-RNF113A expression vectors, then processed for immunofluorescence as shown. **(D)** Quantification of WT and N5 mutant HA-RNF113A from **(C)** for co-localization with the nuclear speckle marker PRP8. N = 3 replicates and error bars indicate ±S.D. of the mean. (**E**) HeLa-S cells expressing the indicated vectors were subjected to RNA immunoprecipitation using anti-Flag resin. The input lysates and the immunoprecipitated material were analyzed by Western blot with indicated antibodies. **(F)** U2OS cells were transduced with the indicated shRNA and Flag expression vectors. Whole cell lysates were used for Western blotting and probed with the antibodies as shown.

