## Supplemental Figures S1-S7 for "Aberrant RNA methylation triggers recruitment of an alkylation repair complex"

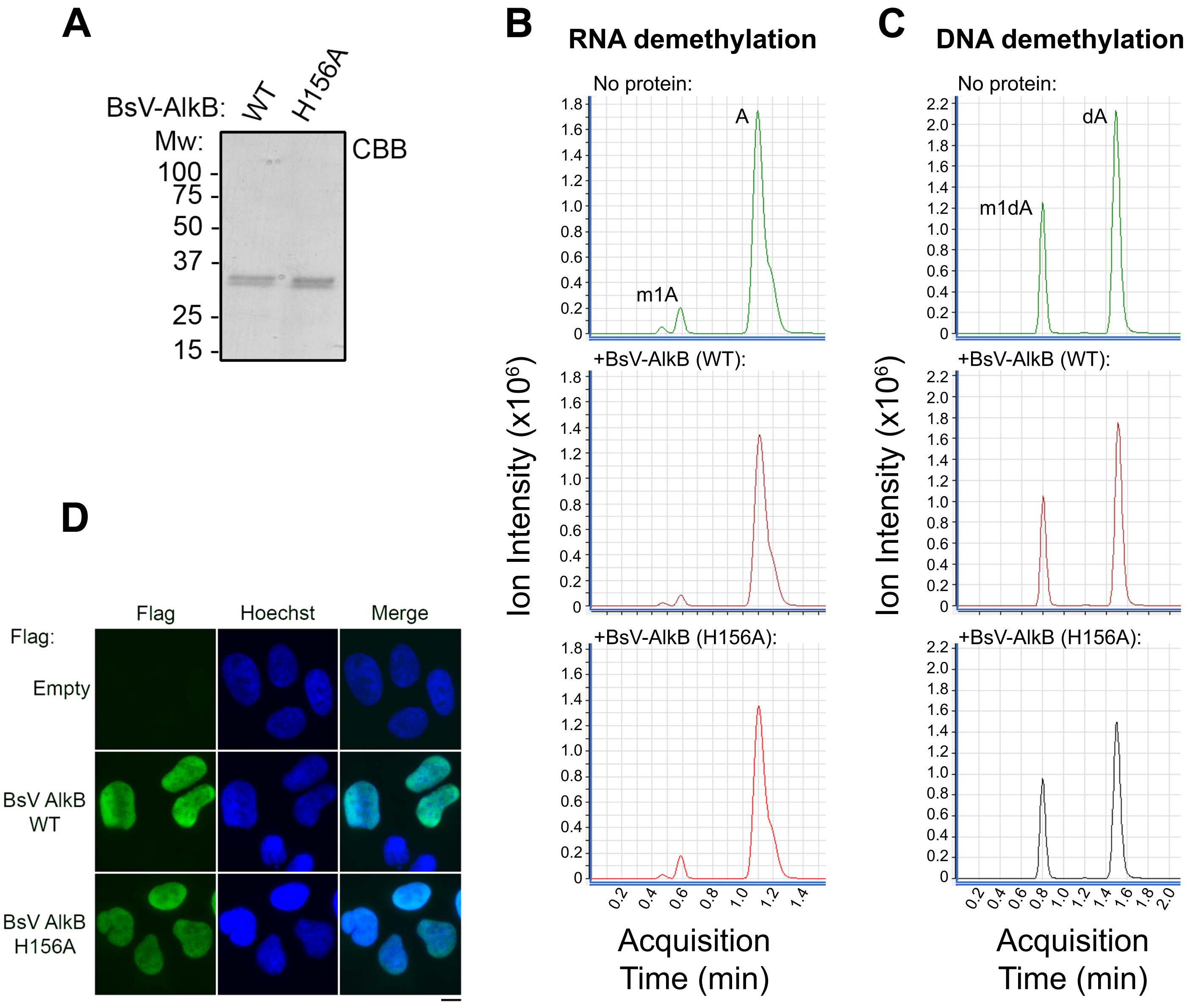
A
BsV-AlkB:
WT
H156A
Mw:
CBB
100
75
50
37
25
15
B
RNA demethylation
No protein:
A
m1A
+BsV-AlkB (WT):
+BsV-AlkB (H156A):
Ion Intensity (x10^6)
Acquisition Time (min)
C
DNA demethylation
No protein:
dA
m1dA
+BsV-AlkB (WT):
+BsV-AlkB (H156A):
Ion Intensity (x10^6)
Acquisition Time (min)
D
Flag
Hoechst
Merge
Flag:
Empty
BsV AlkB WT
BsV AlkB H156A

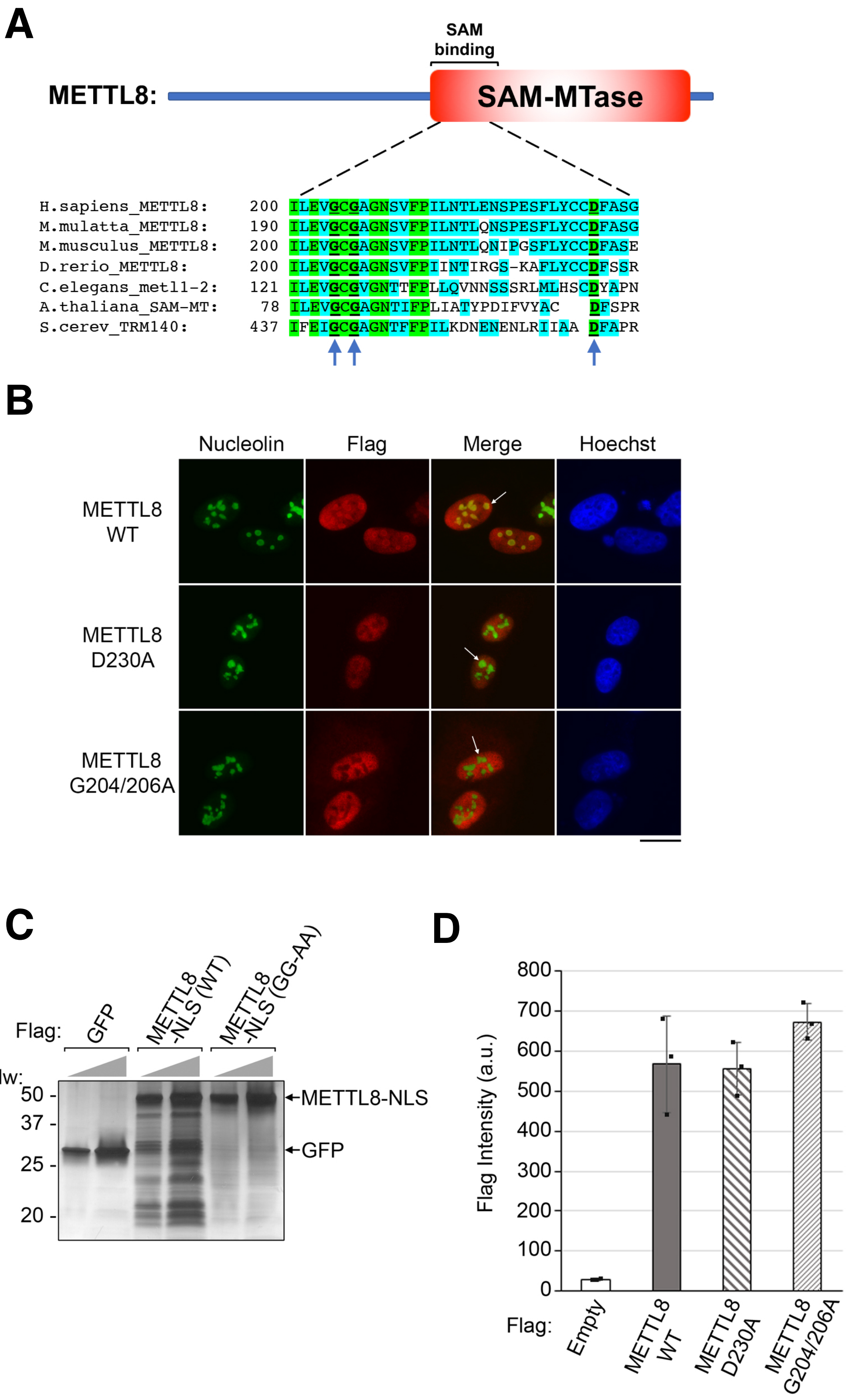
A
SAM binding
METTL8:
SAM-MTase
H.sapiens_METTL8: 200 ILEVGCGAGNSVFPILNTLENSPESFLYCCDFASG
M.mulatta_METTL8: 190 ILEVGCGAGNSVFPILNTLQNSPESFLYCCDFASG
M.musculus_METTL8: 200 ILEVGCGAGNSVFPILNTLQNIPGSFLYCCDFASE
D.rerio_METTL8: 200 ILEVGCGAGNSVFPIINTIRGS-KAFLYCCDFSSR
C.elegans_metl1-2: 121 ILEVGCGVGNTTFPLLQVNNSSSRLMLHSCDYAPN
A.thaliana_SAM-MT: 78 ILEVGCGAGNTIFPLIATYPDIFVYAC DFSPR
S.cerev_TRM140: 437 IFEIGCGAGNTFFPILKDNENENLRIIAA DFAPR
B
Nucleolin
Flag
Merge
Hoechst
METTL8 WT
METTL8 D230A
METTL8 G204/206A
C
Flag:
GFP
METTL8 -NLS (WT)
METTL8 -NLS (GG-AA)
Mw:
50
37
25
20
METTL8-NLS
GFP
D
Flag Intensity (a.u.)
800
700
600
500
400
300
200
100
0
Flag:
Empty
METTL8 WT
METTL8 D230A
METTL8 G204/206A

Figure S3 (Brickner et al.)

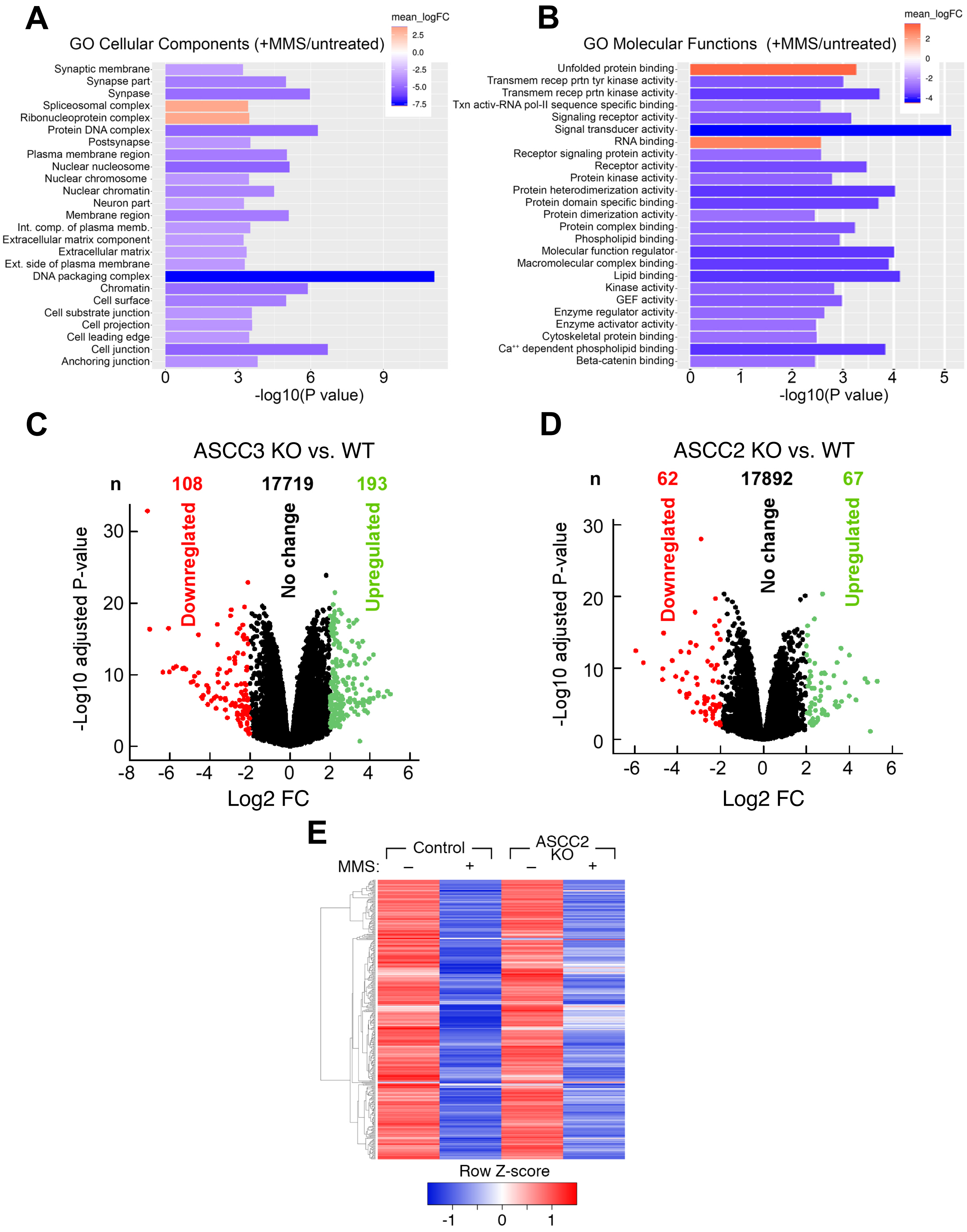

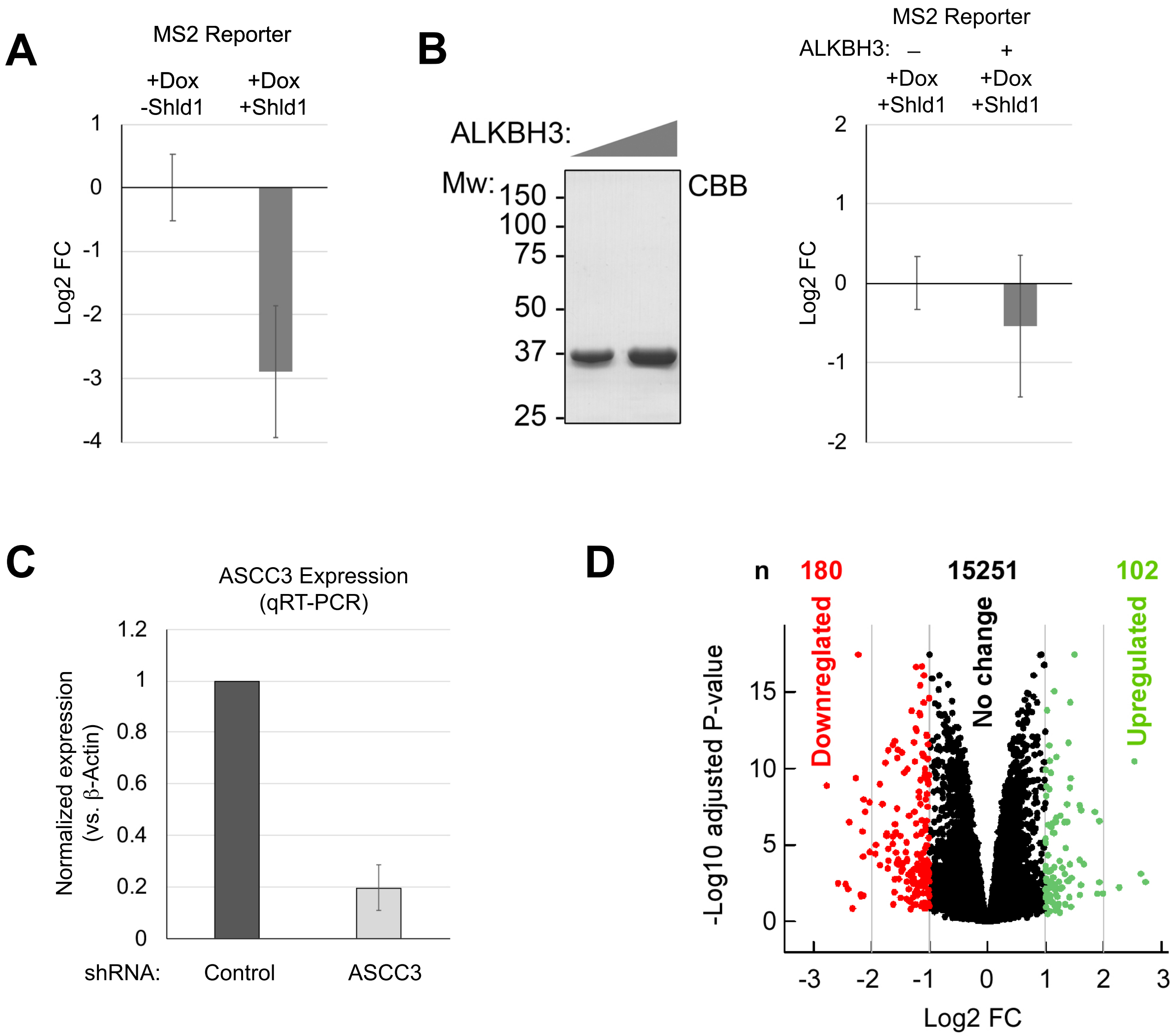
A
MS2 Reporter
+Dox -Shld1
+Dox +Shld1
Log2 FC
1
0
-1
-2
-3
-4
B
ALKBH3:
Mw:
150
100
75
50
37
25
CBB
MS2 Reporter
ALKBH3: – +
+Dox +Shld1
+Dox +Shld1
Log2 FC
2
1
0
-1
-2
C
ASCC3 Expression (qRT-PCR)
Normalized expression (vs. β-Actin)
1.2
1
0.8
0.6
0.4
0.2
0
shRNA:
Control
ASCC3
D
n
180
15251
102
Downregulated
No change
Upregulated
-Log10 adjusted P-value
15
10
5
0
-3
-2
-1
0
1
2
3
Log2 FC

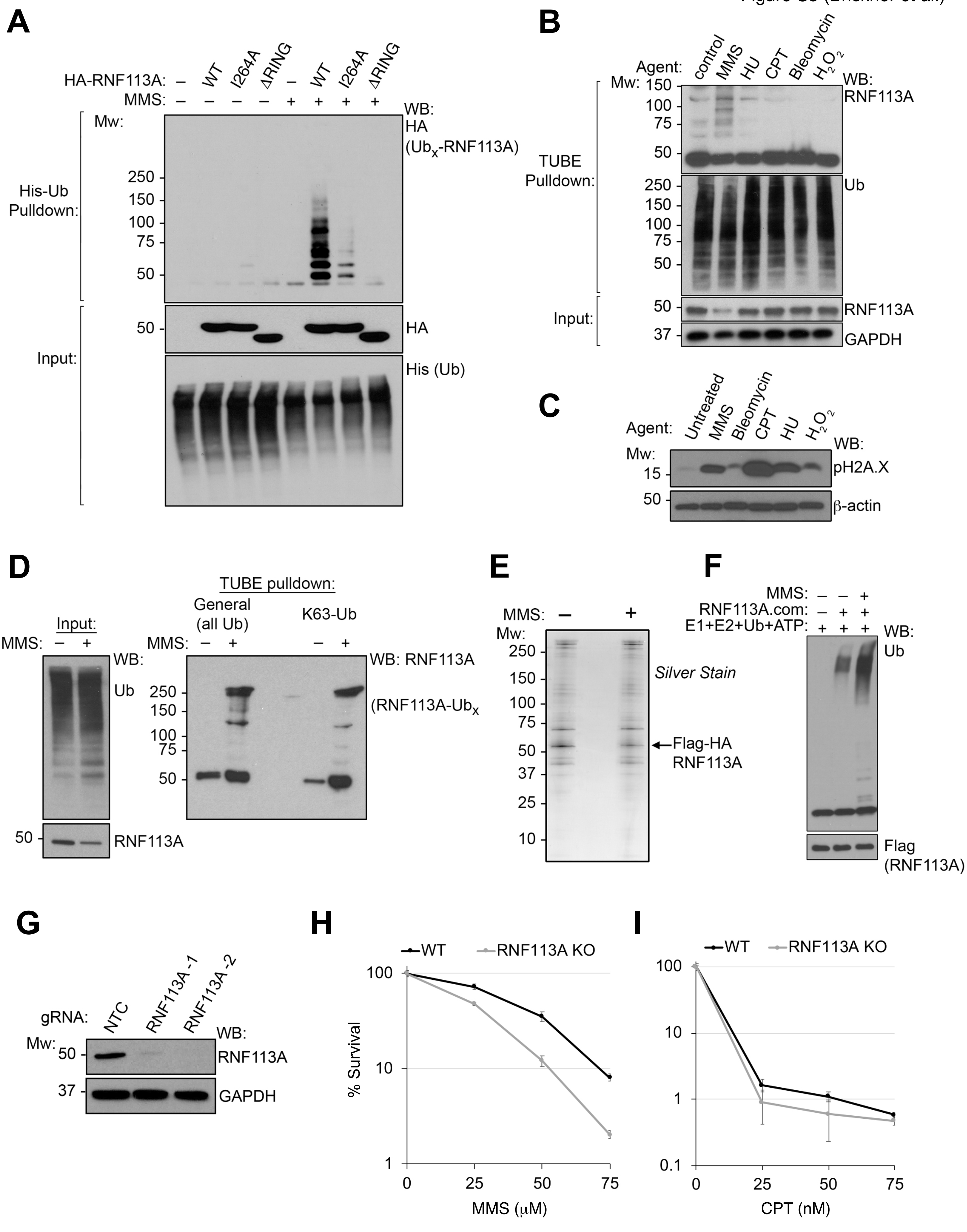
A
HA-RNF113A: – WT I264A ΔRING – WT I264A ΔRING
MMS: – – – – + + + +
WB:
HA
(Ubx-RNF113A)
Mw:
His-Ub Pulldown:
250
150
100
75
50
Input:
50
HA
His (Ub)
B
Agent: control MMS HU CPT Bleomycin H2O2
WB:
Mw: 150
RNF113A
100
75
50
TUBE Pulldown:
250
150
100
75
50
Ub
Input:
50
RNF113A
37
GAPDH
C
Agent: Untreated MMS Bleomycin CPT HU H2O2
WB:
Mw:
15
pH2A.X
50
β-actin
D
Input:
MMS: – +
WB:
Ub
50
RNF113A
TUBE pulldown:
General (all Ub)
K63-Ub
MMS: – + – +
WB: RNF113A
250
150
100
75
50
(RNF113A-Ubx
E
MMS: – +
Mw:
250
150
100
75
50
37
25
10
Silver Stain
Flag-HA RNF113A
F
MMS: – – +
RNF113A.com: – + +
E1+E2+Ub+ATP: + + +
WB:
Ub
Flag (RNF113A)
G
gRNA: NTC RNF113A-1 RNF113A-2
WB:
Mw:
50
RNF113A
37
GAPDH
H
WT
RNF113A KO
% Survival
100
10
1
0
25
50
75
MMS (μM)
I
WT
RNF113A KO
100
10
1
0.1
0
25
50
75
CPT (nM)

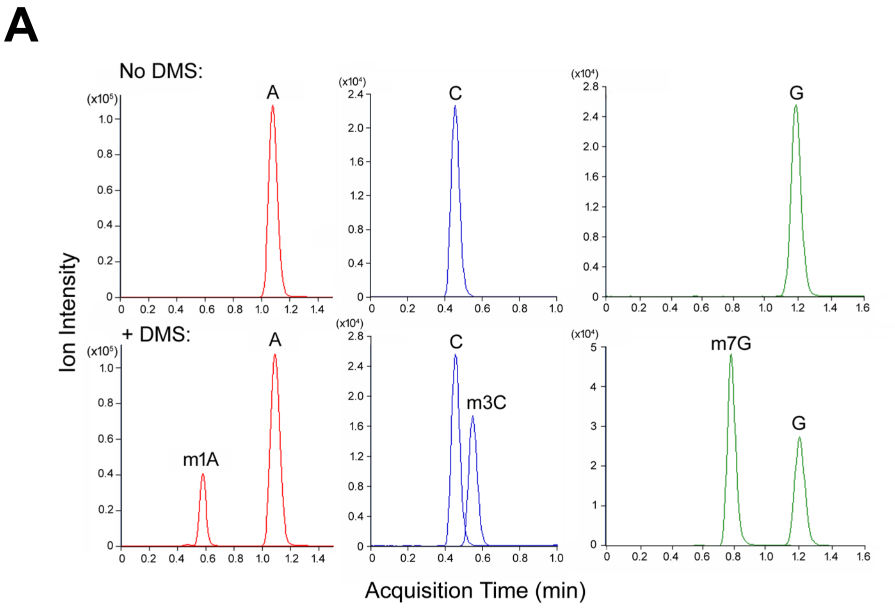
A
No DMS:
+ DMS:
Ion Intensity
A
C
G
m1A
m3C
m7G
Acquisition Time (min)

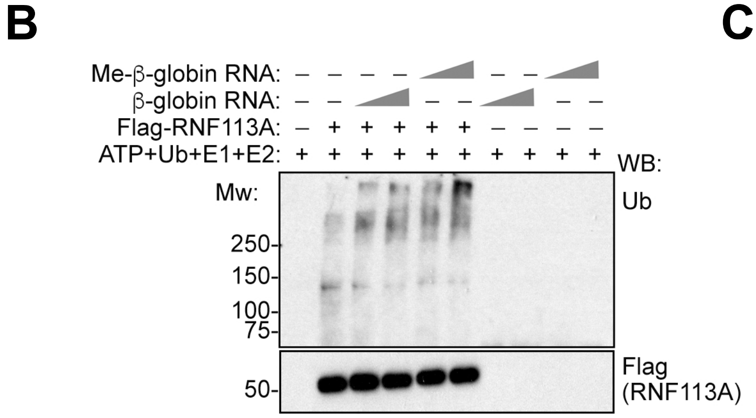
B
Me-β-globin RNA:
β-globin RNA:
Flag-RNF113A:
ATP+Ub+E1+E2:
WB:
Mw:
Ub
250-
150-
100-
75-
50-
Flag (RNF113A)

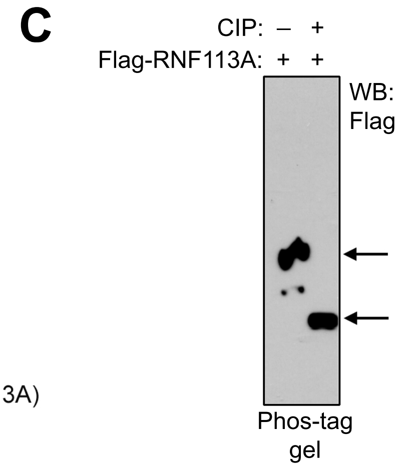
C
CIP: – +
Flag-RNF113A: + +
WB: Flag
Phos-tag gel

A

| Peptide identified by LC-MS: | Phosphorylation site(s): |
|---|---|
| EPIQSTGSMAEQLSPGK | S6 |
| RPACDPEPGESGSSSDEGCTVVRPEK | S46, S47 |
| AAYGDLSSEEEEENEPESLGVVYK | S84, S85 |
| YGVYEDENYEVGSDDEEIPFK | S253 |

B

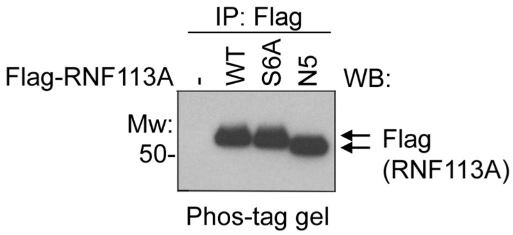


C

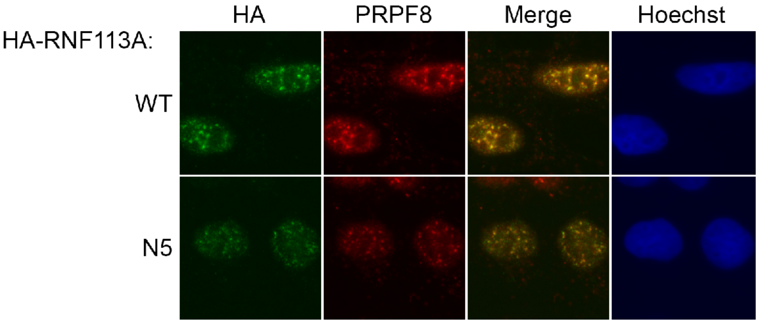


D

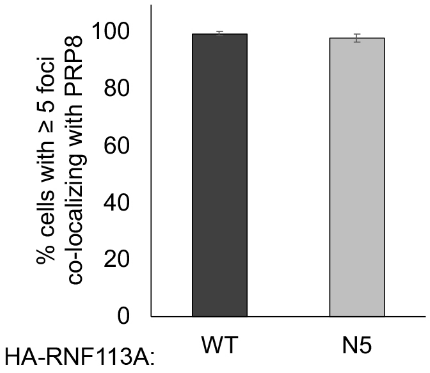


E

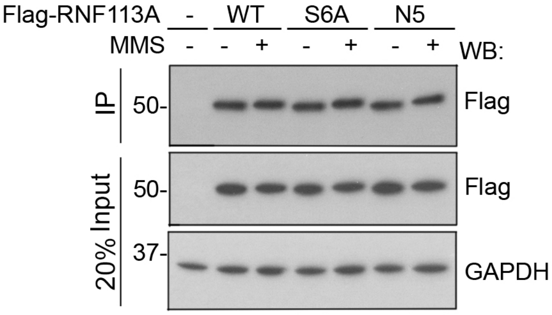


F

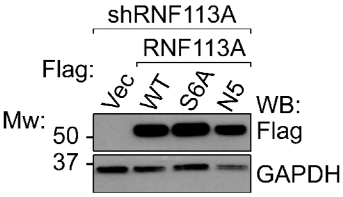
